# Quantifying and modelling non-local information processing of associative brain regions

**DOI:** 10.1101/2019.12.19.883124

**Authors:** Johannes Balkenhol, Juan Prada, Hannelore Ehrenreich, Johannes Grohmann, Jóakim v. Kistowski, Sonja M. Wojcik, Robert Blum, Samuel Kounev, Thomas Dandekar

## Abstract

Brain world representation emerges not by philosophy but from integrating simple followed by more complex actions (due to drives, instincts) with sensory feedback and inputs such as rewards. Our simulation provides this world representation holistically by identical information encoded as holographic wave patterns for all associative cortex regions. Observed circular activation in cell culture experiments provides building blocks from which such an integrative circuit can evolve just by excitation and inhibition transfer to neighbouring neurons. Large-scale grid-computing of the simulation brought no new emergent phenomena but rather linear gains and losses regarding performance. The circuit integrates perceptions and actions. The resulting simulation compares well with data from electrophysiology, visual perception tasks, and oscillations in cortical areas. Non-local, wave-like information processing in the cortex agrees well with EEG observations such as cortical alpha, beta, and gamma oscillations. Non-local information processing has powerful emergent properties, including key features of conscious information processing.

## Introduction

The human brain relies on the complex interplay of different neuronal circuits; both serial processing and neuronal recognition as well as integrative properties and holistic processes are important^1–6^. However, a computer model for this emergent development of such integrative processing is still missing. We find that in our simulation, key features of consciousness emerge including the integration of voluntary movement, a world model and a model of the self^7^. It requires a high enough number of high level perceptrons or microcircuits^7^ in Mountcastle columns to allow serial processing as well as a sufficiently strong non-local wave processing circuit. The simulation suggests that below a critical threshold of 1 billion neurons the integration power of the circuit breaks down and encoded subjective presence becomes too short to allow planning or voluntary action. Anatomical data and various electrophysiological measurements agree well with the *in silico* model. There are matching data on the break-down of integrative functions in Alzheimer’s disease and Schizophrenia. Our high-level simulation provides local processing units with the same information: Mountcastle columns and perceptrons^8^, or basic neuronal circuits such as the gamma generating microcircuits^7^. We provide a highly detailed view on emergence in information processing at different levels (genes, signaling, neurons, brain). Scaling up experiments suggest that no new emergence level arises using grid computing, but instead this produces linear gains in perception and processing as well as performance losses due to communication overhead.

## Results

Netlogo, a multi-agent programmable modelling environment^9^, models signal transmission between neurons, non-local signal processing, neuronal activity, and emergent cortical wave patterns.

Information for survival is (i) selected by evolution and (ii) stored internally on ever higher levels (**Fig. 1a-e**; DNA, encoded protein networks; instructed neuronal networks; human language; **eq. S1**, supplement). Stimulation of signalling cascades and activating neural cells generate wave patterns from simple rules. The BDNF /TrkB protein network illustrates how this is programmed inside neuronal cells on a molecular level^10–14^. (**Fig. 1a-e** and supplement; Video 1, chemical activation of cultured neurons; Video 2, signalled wave patterns in glial cells https://www.biozentrum.uni-wuerzburg.de/bioinfo/computing/neuro). Through such patterns, identical information may be provided to all Mountcastle columns^15,16^, actual decoding or storage happens in individual columns. The resulting central circuit stores, mediates and integrates instincts, drives, and adaptive behavior into emerging conscious thoughts.

**Figure 1:**
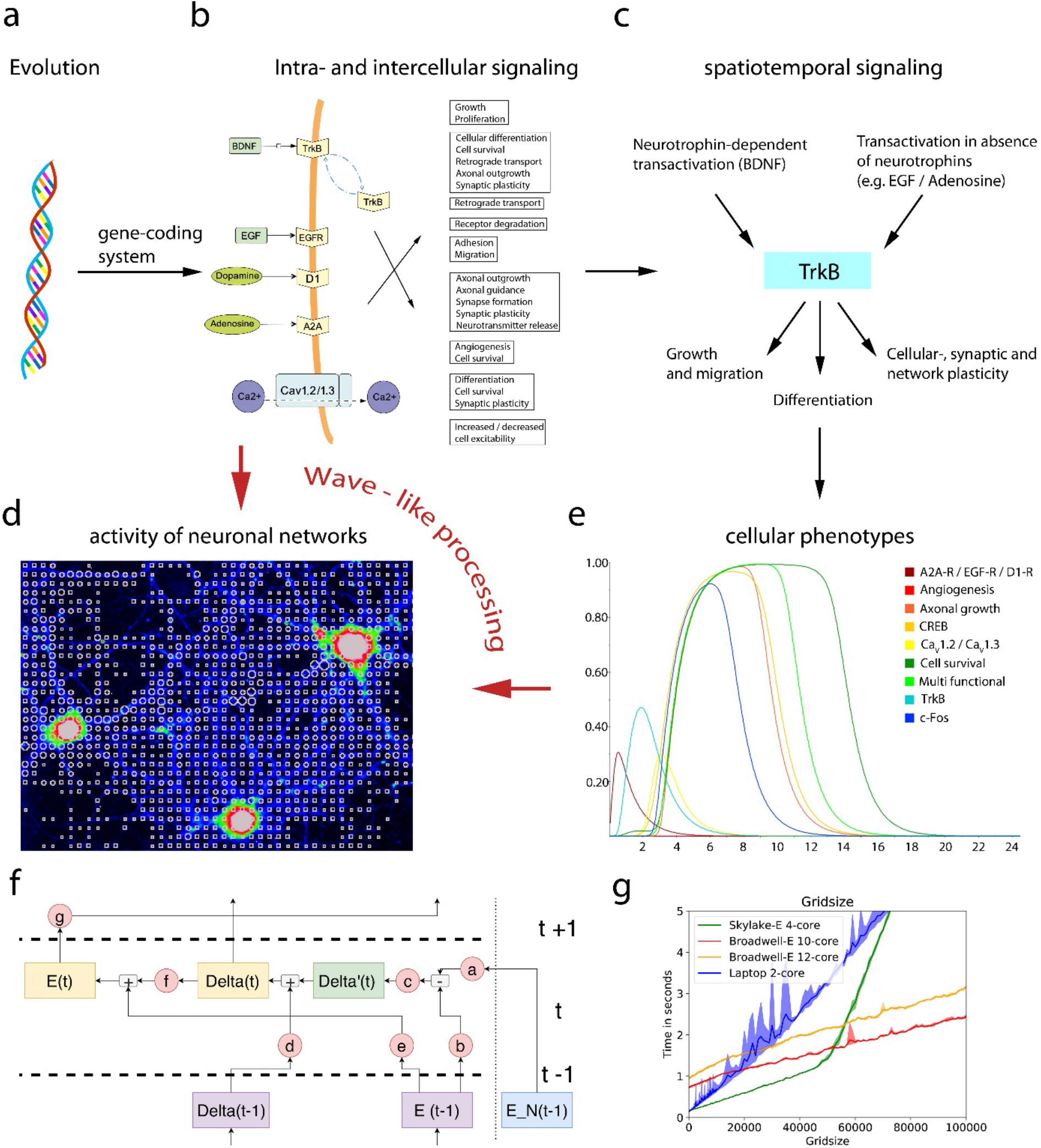
Information is processed on ever higher levels and networks. The principle of selecting better and better adapted information stored internally for survival combined with ever higher levels of nodes and supernodes is sufficient to imply rather complex emergent behavior. Here, different emergent levels of signal processing are exemplarily shown (in a-e) to illustrate how wave-like processing contributes to emergence. **(a)** Genetic information codes the regulation program of cell type-specific signaling networks. **(b)** For instance, signals caused by the brain-derived neurotrophic factor (BDNF) stimulate cellular and synaptic plasticity in cortical regions of the brain by activating the receptor tyrosine kinase TrkB. Cell surface TrkB and intracellular TrkB (two blue arrows, top) are embedded in a complex intra-and intercellular signaling system with other factors (EGF, dopamine, adenosine) to regulate diverse functions in many cell types (details: supplement). **(c)** For this specific protein network, spatiotemporal activation of diverse cascades is caused by neurotrophin-dependent effects through BDNF/TrkB or by transactivation of TrkB in the absence of neurotrophins. TrkB activation in the absence of neurotrophins can cause migration, differentiation, or survival of neural cells, e.g., early in development, but can also affect differentiation and plasticity in the adult brain. **(e,** right) However, a new emergent effect can arise if the signaling interactions happen between cells, for instance if differentiation of cell types is signaled. Exemplarily, this simulation shows how neurotrophin-independent activation of TrkB by either adenosine receptor A2A-R, EGF-receptor (EGF-R), or dopamine receptor D1 (D1-R) activates specific key nodes (e.g. the transcription factor CREB) to participate in different cellular functions, such as information processing. Other functions (e.g. angiogenesis) are not modelled by this Jimena software simulation (x-axis: activation state; y-axis: time, arbitrary units). Modulating the decay time after TrkB activation can block the neurotrophin-independent contributions to this basic level of emergence in cellular communication (details: supplement). A biological explanation of this simulation might be the regulated cell surface abundance of TrkB. **(d,** left) With synaptic communication between neurons, new dynamic properties of a neuronal network, for instance synchronized neuronal activity with wave-like properties, can emerge. This phenomenon is indicated here by chemical activation of hippocampal neurons in culture (Video 1; https://www.biozentrum.uni-wuerzburg.de/bioinfo/computing/neuro). The image shows loci of calcium activity (white circles) projected onto an average-projection image of hippocampal neurons loaded with a fluorescent calcium indicator after stimulation with a chemical LTP solution (modified from^45^). Fluorescence intensities are indicated by a rainbow look-up table. Similar wave-like patterns are also observed in glia cells showing long-range, wave-like signals after local stimulation with glutamate (Video 2, link above). **(f)** Such wave-like patterns for non-local information processing rely on **holographic brain microcircuits**. To simulate this, three processing steps of each node build on each other (see eq. 1 and Figure 2c): First, the derivation of the neighbouring energy (*E_N(t-1); activation1n(t*)) and the own energy state *(E(t-1); activation1(t)*) leads to the *Delta’(t)* value (*slope_vector(t*)) of that specific node. Second, this *Delta’(t)* value is integrated with the *Delta(t-1)* function (*slope_old*(*t-1*)). Third, the updated *Delta(t) (slope_old(t*)) is added to the energy state function of the focused node (*E(t-1);activation1(t*)) leading to the current final energy state (*E(t);activation0(t)*). Both the new delta (*Delta(t); slope_old(t*)) and the new energy state (*E(t);activation0(t*)) are then forwarded to the next time step, while the current energy state (*E(t-1);activation1(t*)) is forwarded to all neighbouring nodes. The variables *a* (*ratio_neighbour_activation), b (ratio_inhibition_activation1), c (neighbour_integration; NI_slopev), d, e, f (slopeo_damping*) and g (*damping*) are coupling factors. These modulate the energy transfer between the neuronal nodes. Their default value is 1. The variables can be replaced by a function e.g. a damping function (details in supplement). **(g) computer cluster simulation of non-local information processing in the brain.** Shown is a perception task model with 5000 ticks with a resolution of 1 tick per millisecond (i.e., 5 seconds of real-time) and its dependence on the number of simulated nodes in the cluster. The perception task is done more efficiently with more nodes but there are no emergent new properties. Parameters: Average run-time of 5 repetitions are shown as lines, the minimum and the maximum run-time as semi-opaque areas of the same color. We are able to simulate up to 50 000 nodes in real-time (i.e., the simulation runs faster than the modelled process itself) using a standard 2-core laptop. However, it is possible to simulate even more nodes (100 000 and higher) by running the simulation on server CPUs. We furthermore observe that the servers show less random influences compared to the laptop workstation. This is expected, as we used bare-metal servers with no other competing processes running at the time of the simulation. Additionally, we observe that both the laptop and the smaller Skylake server are faster for smaller simulations, but are strongly influenced by the number of nodes to simulate. The reason for this is the smaller L2-cache. In fact, we observe a strong impact on the performance of the Skylake server at around 500000 nodes. This is exactly when the node grid gets too big for the L2-cache, which has a serious impact on performance.

This central circuit displays key properties of consciousness, including the integration of sensory inputs with voluntary actions. It results from basic neuronal circuits of shells of activating and inhibitory neurons (about 10^4^ Mountcastle columns or 10^9^ neurons). A world representation iteratively evolves from such a central circuit not as a philosophical insight but rather from a feedback loop of simple and then ever more complex actions and resulting rewards. Encoded as a holographic wave pattern, the same information reaches all conscious cortex regions: the same world view, and it integrates actions (“will”).

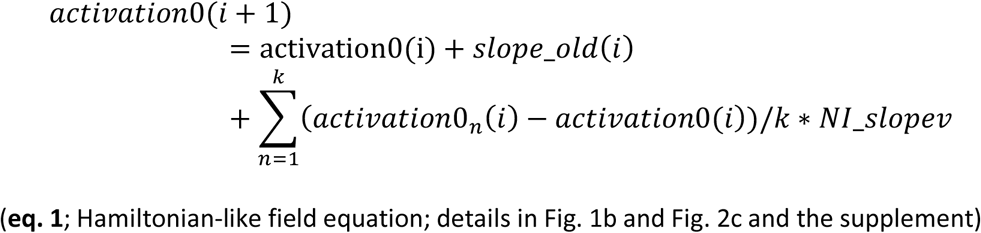

Next, a high number of our simulations (**Fig. 1a-e and f**) were coupled on a large computer grid (opensource C code available at https://github.com/DescartesResearch/BrainSimulation). Parallel computing sped up the perception simulations (**Fig. 1g**), exhibiting constant performance due to the linear increase of computation time based on the number of nodes. We simulated up to 400 000 nodes in real time using a resolution of 1 tick per millisecond and 24 core server system with 6 MB of cache (**Fig. 1g**). Perception speed as well as communication and processing time all increased linearly. Hence, after emergence of a holographic medium (simulating 10^4^ Mountcastle columns of 10^5^ neurons each for decoding)^155^, further processing power did not lead to new emergence levels but just to linear gains and losses.

In detail, diverse sensory input is transformed into holographic interference patterns (**Fig. 2a**; three external and two proprioceptive signals). The same information reaches all pixels, individual Mountcastle column in the simulation. The wave pattern results from symmetrical interaction of all neighbours, a grid of 100 × 100 =10^4^ Mountcastle columns, about 10^9^ neurons in total, participating in the simulation. This mirrors basic qualities of consciousness, transforming diverse signals into a unified world view stored in a central wave pattern shared by all participating Mountcastle columns. The interference pattern can easily be decoded: **Fig. 2b** shows different waves involved and overtones. The storage capacity of 3x 50 bits during 3 seconds also corresponds quantitatively to conscious perception. Proprioceptive signals integrated into the circuit include motor action and pain signals, external world input covers all sensory qualities to build up a representation of self and world^17^ using new sensory input as feedback (**Fig. 2a**). **Figure 2c** presents a typical neuronal excitatory and inhibitory microcircuit^7^ that mirrors the algorithm described in **Fig. 1f** and **eq. 1**. Thus, a substantial amount of neuronal brain activity is found to be non-local and wave-like^18–20^, instead of direct, computer-like logical connectivity. We, therefore, hypothesize that the conscious, associative regions of the brain are the neurobiological analogue to the algorithm.

**Figure 2:**
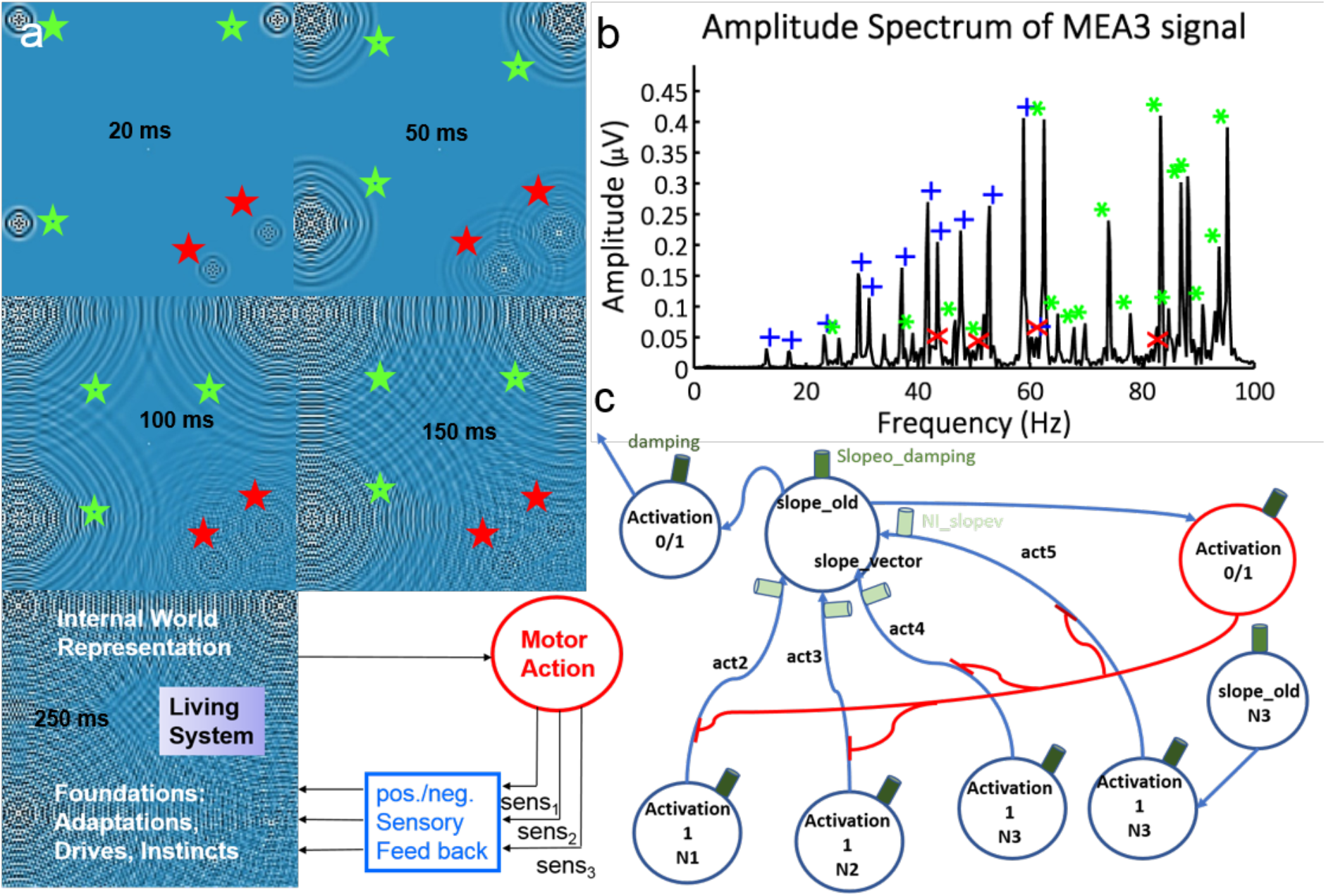
A neuronal circuit for non-local information processing. (a) Wave patterns are formed by symmetrical interaction of neighbours. The internal and external stimuli are copied, statistically processed, compared to existing information, and integrated. As a consequence, the entered information is distributed over the whole model, which makes it available at any column after a distinct time period. Red stars indicate external input, while blue stars indicate internal input. The pictures were made at 20, 50, 100, 150and 250 ms after stimulus onset. Processing of information is time dependent. Hence, the information spreads over the whole model founding the basis for an experienced presence (here ~200 ms). Fully integrated information can be depicted from each location. The information can be coded in harmonics and phase by means of the complex interference pattern. With an FFT, the input information can be decoded. The frequency coding is dependent on sampling rate, wave speed and wavelength. This results in an input/ output system of a simulated brain slice exemplified as a wave interference model with emergent world and self-representation properties. Input from the external world coded in frequency and phase is integrated into an emergent system that consists of unified processing units. The overall model is a live processing system. In a living system there is permanent feedback to actions caused by drives, instincts etc. by input from the external world. An internal representation of self and outside world results^44^. **(b) Decoding: Fast Fourier transform-analysis (FFT) of the neuronal circuit.** The example shows how, at any place of the simulation, for instance in the multi-electrode array position 3 (MEA3) decoding can be efficiently done. The decoding shown is an electrode signal of 12 input frequencies (blue) and generated overtones (green) that appear when periodic spike input is applied. Red crosses indicate noise. Only one electrode is presented. **(c) The holographic brain circuit allows processing calculations, high discrimination and signal resolution** (compare eq. 1 and Fig. 1b). The 4 neighbours (N1, N2, N3, N4), transfer their energy to a representative neuron of a central processing unit. The currently calculated activation of the central processing unit (*activation1*) is comparing its own energy level to that of each neighbouring activation, means the *activation1* is subtracted from each *activation N1-4* (red lines). The subtraction is mediated by inhibition on axonal level. This results in *act2, act3, act4* and *act5*. Together they are integrated as *slope_vector* and added to the former *slope_old* old value of a representative integrating neuron. The energy level of *slope_old* is transferred and integrated into the former activation level of *activation0*. From there on *activation1* is updated and *activation0* itself contributes to the update of energy level of its neighbours. Via different types of ion channels mediating phasic (synaptic), tonic (cell body inhibition, or axodendritic inhibition) the signal transfer within the neuronal circuit can be influenced. The coupling factors *neighbour_integration (NI_slopev*) determine the extent of calculated *slopev*, whereas *slopeo_damping* and *damping* determine the extent of updated *slope_old* and *activation0* (details in supplement).

Our non-local, wave-like and holographic processing simulation is highly compatible with EEG and MEA recordings (**Fig. 3 and 4**). This holistic system state perspective indicates possible target points for influencing physiological coupling parameters resulting in alternating systems states, such as sleep, waking state, anaesthesia, and schizophrenia (for a detailed description, see **Fig. 2c**, **eq. 1**, and supplement). Peak distributions of multiple electrode arrays (MEA)^21^ underlying the EEG pattern of slow wave sleep, awake state (freely behaving) (**Fig. 3a**), and anaesthesia network state compare well to our simulation (**Fig. 3b,c**). Ribeiro *et al*.^21^ show an equivalent lognormal tail distribution of awake and slow wave states, as well as the transformation to power law distribution with onset of anaesthesia. Additionally, detailed parallel analysis of *in silico* (**Fig. 3b**) and *in vitro* electrode (**Fig.** recordings **S9a,b**) show evident similarities in lognormal distribution using the same analytic method. The hippocampus^22^, as well as the model, produce overtones whenever peak input is applied (**Fig. 3d**). Using sine input, the overtone generation is decreased and nearly inhibited in the hippocampus^22^ which is also valid for the simulation (**Fig. S2**). There is a linear dependence of the speed at which the waves travel and the maximum frequency that is found in electromagnetic measurements^18,19^ and in the model (**Fig. 3e**). This collapses at maximum speed of 0.5 m/s and frequency (500 Hz), the maximum sampling rate of the model.

**Figure 3:**
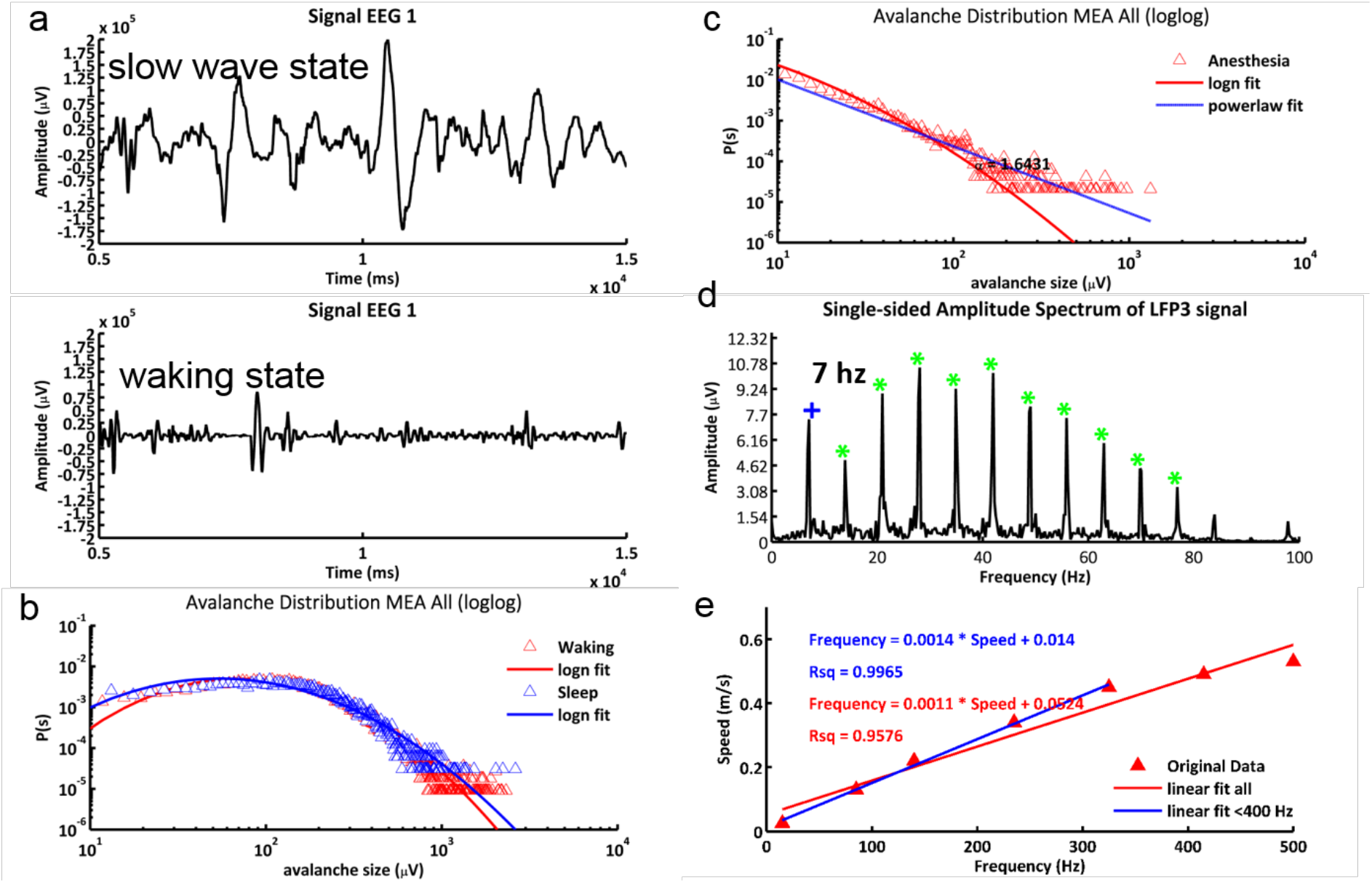
Model validation by EEG data. (a) reproduction of EEG asleep state (slow wave activity, top); and waking state (below). References showing highly similar experimental data are given for each subpanel. Top shows increased amplitude while the bottom signal shows a higher frequency. A low pass filter and can be used to model the slow wave state (decrease of *NI_slopev* to 0.1), as well as a slight decrease of the carrier frequency (0-13Hz) and *slopeo_damping* (1.0E-3), as well as damping (1.0E-5). Transfer to waking (freely behaving) urges increase of *NI_slopev* to 2.6655 along with slight increase of *slopeo_damping* (1.0E-2), *damping* (1.0E-4), and carrier (0-23Hz). The size distribution of positive peaks reproduces MEA peak distribution in **(b) freely behaving** (sleep and waking) and **(c) anesthetized state**. Freely behaving state: lognormal distribution. Observation data: Ribeiro *et al*.^21^. A power law distribution fits better for anesthesized rats (*damping* and *slopeo_damping* is increased to 0.001 and 0.1, whereas *NI_slopev* stays at 2.6655). **(d) Systematic analysis of overtones** generated by the simulation. The cortical architecture of the brain (rodent hippocampus) can produce similar non-local overtones^22^ from short pulsed peak input, sinusoid input decreases these (compare to data in Laxpati *et al*.^22^, Figure 7). **(e) Wave speed** increases linearly with maximum frequency in the model. Via the *NI* energy transfer the maximum frequency is adjusted. The speed at which the wave hills travel decreases linear with decrease of the energy transfer (*NI_slopev*). With decrease of *NI_slopev* the maximum frequency that can be generated and decoded is decreasing linearly. Typically, due to the overtone generation the maximum frequency is limited by the highest overtone. Close to 500 Hz and an *NI_slopev* of >2.6, the speed of the simulated waves converges to 0.5 m/s. A wave speed of cortical waves of 80-107 mm/s at a frequency of ~7 hz is determined by Lubenov and Siapas (Figure 4)^18^ in the hippocampus. Furthermore, a linear relationship between wave speed and frequency is described by Zhang and Jacobs 2015 (Figure 4)^19^.

As estimated for conscious perception^23^ and subjective presence^24^, the simulation easily encodes and stores over 50 bits of information within 3 seconds (**Fig. 4a**). The half-width of the peaks using sinus wave input is 0.5 Hz within 1 second for this model (see **Fig. S6**). With a bandwidth of 10-110 Hz, even 200 bit/second could be coded. Within these ranges, model size has no influence on the decoding of the input frequencies and low influence on the generation of overtones (**Fig. 4b)**. As simulated (**Fig. 4c**), spontaneous baseline activity of cortical areas organizes after several seconds to minutes without external stimuli^25,26^ around ~8 Hz^27–29^. Brain coherence measurements without external stimuli^30,31^ align well with the coherence patterns of simulated spontaneous activity in dependence of space and frequency (**Fig. 4d**). The simulated power law behaviour of degree distribution for EEG or FMRI fits EEG or FMRI observed (**Fig. 4e**) and an Ising model of brain criticality^32^ including variations to altered average connectivity (value k) (**Fig. 4e**). Different pathologies can easily be studied in our model, e.g., effects of multiple lesions during Alzheimer’s disease degrading information processing (**Fig. 4f)** and the lack of neuronal correlation on coding in Schizophrenia with a subsequent shift of real input frequencies to non-coherence (**Fig. 4g**). Moreover, a self-organizing theta and beta-band following external stimuli superimposed on a holographic carrier was generated in our model and we simulated the beta band decline in Schizophrenia (**Fig. 4h**). Both effects notably match what has been described in the literature^33,34^. Fast ripples^35^ arise during such a beta band generating stimulus period (**Fig. S8**). Based on the integrated analysis and the effect of undersampling (**Fig.S10**) we suggest that information in the cortex is encoded at a bandwidth >400 Hz. High frequency coding is well supported^36^.

**Figure 4:**
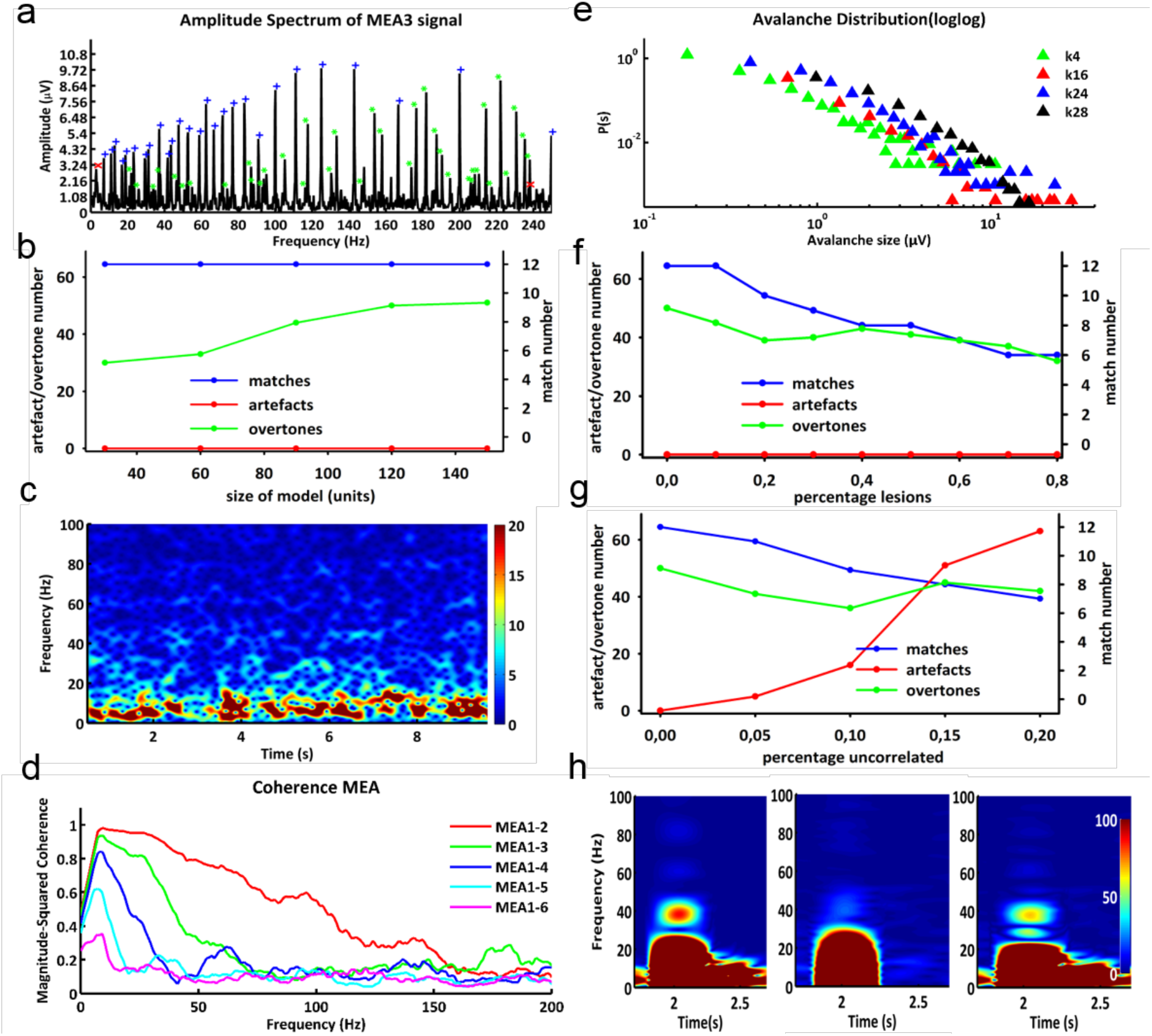
Properties of non-local processing of stimuli in a holographic brain model. (a) Efficient resolution of 28 different stimuli as complex model input. The input is coded in prime numbers as waves lasting from 5 to 239 ms. The corresponding frequencies are 5 - 250 Hz. 28 out of 29 input signals and 45 overtones could be decoded. For comparison, the undetected frequency is in the non-coding space (<7 hz). **(b) Scalability of model-size: similar wave separation power.** A 20×20 matrix, as well as a150×150 matrix is equally resolving 12 peak stimuli (13, 17, 23, 29, 31, 37, 41, 43, 47, 53, 59, 61 Hz) without generating artificial frequencies. The matched frequencies, overtones, and artificial frequencies are investigated. The x-axis represents the diameter of processing units. The left y-axis corresponds to the number of detected artificial signals and overtones. The right y-axis represents the number of input frequencies. **(c) Self-organizing baseline activity of the simulation mimic observations.** This is generated by applying solely spontaneous activity of 100 mV and 0.01% of activated processing units each time step in waking state. **(d) Spontaneous activity and coherence of human brain mirrored.** MEA coherence measurement of spontaneous activity (0.01% activated cells firing with 100 mV) in absence of stimuli. The stimulating signal lasts 10 seconds. The distance of the MEA electrodes is 0.5 (MEA1-2), 2 (MEA1-2), 4 (MEA1-3), 8 (MEA1-4), 16 (MEA1-5), 32 mm (MEA6). **(e) Avalanche distribution and criticality correspond to the human brain.** The size distribution of positive peaks in the EEG signal is analyzed according to the number of neighbours building a directly connected processing unit (k value). Parameters are the same as in Figure 3a (waking), but the number of neighbours is altered. **(f) Influence of lesion occurring during Alzheimer’s disease on information processing.** Axis and parameters are similar to Fig. 4b, except the model size is fixed (150×150 processing units) and the x-axis represents the ratio of lesions. **(g) Low correlation such as in Schizophrenia decreases information content and input separation.** Axis and parameters are as in 4f. The correlation of columns is altered, as indicated by the x-axis. **(h) Beta band decline in schizophrenia.** The self-organizing beta band following a stimulus superimposed on a carrier is shown for healthy (left) subjects and Schizophrenic subjects (middle). (right) is the delta of (left) and (middle). The parameters are the same as in 3a (waking), whereas the carrier is fixed to 10 Hz and the stimulus to 430 Hz. Th burst length is 30 ms. Here (middle), the correlation of information transfer decreased. The color bar indicates activity in μV.

Large-scale models of the brain include the laminar cortex architecture^37^ and detailed neuroanatomic models^38^. We focus on holistic function and properties of consciousness emerging from non-local information processing where holographic models are known to provide a strong foundation^1,2,5^. However, we provide here a first detailed simulation and quantitative model. Validation data presented include direct observation of basic neuronal circuits, data from hippocampal brain slices, as well as EEG und functional data from animal experiments and a human brain. All support our simulation of substantial non-local, wave-like information processing in the cortex. This includes (Fig. 3, 4 and S2-S7) slow wave and waking oscillations as self-organized waves corresponding to sleep and waking state^21^ and electrophysiological phenomena: existence of spontaneous baseline activity^25–29^, occurrence and decline of overtones^22^, (de)coherence behaviour^30,31^, slow wave and waking signals^21,39^, related avalanche distribution in freely behaving and anesthesia state^21^, wave speed behavior^18,19^, self-organized theta and beta band formation after stimuli^33,34^, avalanche distribution in face of connectivity^32^, information loss during A.D.^40–42Ref^, as well as decoherence^33,34^ and shifted decoding in Schizophrenia^43^. The central circuit simulation has high discrimination and signal resolution (**Fig. 4a**). Detailed parameters (**eq. 1**; see methods and suppl. material) fit well to known physiology; alternative settings for pathologies and non-physiological ranges were also tested.

The approach integrates **different key theories of consciousness** (see also supplement): It implements Tononi’s model^3^ of integrative information processing, incorporates Uexküll’s world representation by sensorimotor feedback^44^ and Bohm’s holonomic brain theory^1^ hereby is linked with a concrete circuit and model. Consciousness is probably a real phenomenon, an emergent property of non-local information processing from wave patterns already present in smaller brains. With a critical number of neurons (about 10^8^), a holographic medium results, integrating conscious motor, proprioceptive and sensory input into a unified model of self and world representation. With a higher number (about 10^9^), it integrates at least 150 bits for several seconds allowing to plan and reflect (at least 3 seconds in simulation, as observed^24^). This subjective presence is provided as a holographic brain pattern to all participating Mountcastle columns.

For the first time an open-source simulation encapsulates non-local information processing and decoding. Resulting shared wave interference patterns allow to provide the same information to all participating cortex regions. This may be sufficient to allow emergence of consciousness. While it is too early to unequivocally confirm this notion, our simulation reproduces all available observational data, including detailed brain electrophysiological wave patterns.

## Supporting information

simulation details

## Methods

Our model predicts the emergence of higher neuronal functions from DNA encoded wiring of networks (level 1) and then suitable neuronal circuits (level 2) such that non-local processing of information results, allowing on level 3 the emergence of neuronal circuits and the neuronal transport equation (eq. 1 in results).

### Molecular Networks (level 1)

(i) Storage of internal information in the form of genes as well as all higher levels of information storage are central to *life*: A master equation (**eq. S1**)^46,47^ is introduced to calculate increase and decrease of different DNA species or higher levels of information storage in more complex organisms. (ii) Higher levels of information storage also allow for neurons, brains, and learning. Only with this higher neuronal level arises an organism-specific “*meaning*” in the form of adaptive behaviour and understanding of the environment (with multiple and ever more opportunities to identify the meaning of all and everything in the best way to improve survival).

### Cellular model (level 2): BDNF network

The model presented in **Fig. 1b** and in detail in **Fig. S1a** (Topology) and **Table S1** (all nodes, interactions, references) were assembled starting from the KEGG pathway “ko04722 Neurotrophin signaling pathway”^48,49^ and editing it further. This pathway presents the direct activation mechanisms of the neurotrophins NGF, BDNF, NT3, and NT4 and the subsequent activation of the intracellular signalling cascades including the MAPK, PI-3, kinase and PLC pathway. The activation of these cascades results in functions like axonal growth, cell migration, cell survival, cellular differentiation, and plasticity among others (**Fig. S1b**).

### Neurobiological Model - Holographic non-local circuit (Netlogosimulation)

For the purpose of setting up a brain slide simulation, Netlogo 6.0.4, a multi-agent programmable environment, is used^9^. This is a combined approach of programming environmental and visual representation, as well as graphical modulation. The program is available at https://www.biozentrum.uni-wuerzburg.de/bioinfo/computing/neuro, further simulation details are explained in the supplement.

The simulation models a cortex area of 60 mm in diameter. The input is coming from a lower brain centre, like the thalamus, which is implemented by a carrier signal. In this model, the input entering the cortex area distributes over the cortex area composed of neuronal columns-like integrating units. The information between the units is processed according to the non-local information processing hypothesis. This means the information decoded in frequency and amplitude, reaching the cortex is shared between all neurons in a wave-pattern like manner. As every unit is processing the information and forwarding it to its neighbours in a similar way, according to the unified neocortical column model^50^, the graphical representation appears wave-like. In consequence, information can be depicted for every unit of the model. As the wave-like distribution of the information can explain the appearance of the wave-pattern measured by EEG-electrodes and MEA-electrodes, the output is captured by simulating MEA-electrodes and EEG-electrodes. The summation of a local population of neurons measured by an MEA-electrode is similar to a local-field potential^21,51–53^. According to the EEG-electrodes, the signal is composed of an approximated number of around 1300 columns considering an EEG-electrode radius of around 10 mm^54,55^ and a neocortical column diameter of 0.5 mm. The interelectrode distance of the MEA is dynamically adjusted to the model size and is 10 mm for this study if not stated differently. Further, the signal, which is measured by each electrode of the MEA, is likely to examine the activation level of a single activity of one neocortical column (diameter = 0.5 mm). The signals measured by the electrodes are displayed in 12 plots, 6 MEA-plots and 6 EEG-plots, plotting time against activation level. The activation levels are stated in μV for the MEA and for the EEG. Moreover, the model represents a cortex area of 14 400 neocortical columns. Due to the hypothesis that neocortical columns consist of units of around 10 000 neurons^50,56–58^, the model would come up to 144 000 000 neurons. This is close to the number of columns approximated for the visual cortex of 20 000 columns and the somatosensory cortex of 5 000 columns, with respect to their size of 77 mm and 24 mm in diameter^59,60^. Since the integration of several inputs is creating a complex interference pattern, the information storage capacity of frequency and amplitude is analysed with a fast-fourier transform and a peak-distribution approach described by Ribeiro^21^. The analysed bandwidth is 0–500 Hz. The analysis is made by Matlab 8.2.0.701, a high-level interpreted language, primarily intended for numerical computations (MathWorks, http://www.mathworks.de/products/matlab/).

The power of the model is dependent on, e.g., the size of the model, the frequency restrictions (bandwidth), the number of interacting neighbours, the energy transfer between the neighbours, the influence of inhibitory neurons, the damping, the properties of the margin and the intensity and synchrony of input. Our simulation of non-local information processing can be extended by several modification possibilities. The simulation can be modified by alternating the strength of the signal input and of the carrier signal. In order to investigate the influence of synchrony, the number and strength of neurons activated in the input region can be varied. Further, the coordinates of the input region are variable and can be synchronized. The frequency of the input signal can be individually adapted for each of >=21 signals. In addition, the signal strength of each input can be modulated individually. The information transfer is dependent on the margin damping, the general influence of slope of energy transfer, and the individual influence on the slope of energy transfer of neighbouring columns. There is also a wave direction approach included that enables the user to modulate the energy transfer towards favoured traveling direction of the wave front. As spontaneous activity is described as a distinct feature of the telencephalon^61^, it can be activated and adjusted in ratio and activation strength. Furthermore, the simulation of Alzheimer’s disease and Schizophrenia are added to the model.

The simulation parallels electrophysiological phenomena, e.g., coherence decline, self-organized baseline formation, overtones, slow wave and waking signals, avalanche distribution in freely behaving and anesthesia state, wave speed behavior, self-organized theta band and beta band formation when a stimulus is applied, avalanche distribution in face of connectivity, information loss during A.D. and Schizophrenia and formation of artificial signals production in Schizophrenia, existence of spontaneous activity and cortical size influence.

Further, paralleling all those electrophysiological phenomena *vice versa* leads to the discrimination, resolution, adjustment, decoding, and determination of biologically relevant parameters that underlie brain states and cortical processing (compare Figure 2c).

On top of this, the simulation offers the possibility to extrapolate by tuning the parameters at levels that would be physiologically either devastating or simply not reachable in nature. Doing this, allows furthermore detailed comparison to physical systems, such as different holographic processes, for instance, holograms and holographic storage of image information. Additionally, brain physiology is here turned in to a mathematical formalism (**eq. 1**) and made comparable in a semantic way to other systems treated similarly. The simulation may even suggest a foundation for other critical equations, due to emergence, e.g. sinus-like waves with operating frequency bandwidth originating out of chaos for any type of non-local information processing (**Fig. 4c; S3, S4**).

All results presented are closely paralleled by existing experimental studies, as cited evaluating cortical phenomena.

### Mathematical descriptions

The non-local information processing hypothesis suggests initial uniformity in processing and morphology of each unit. The information entering the processing system is distributed system-wide. The processing of the model is discrete in time. The energy change of each time step of each unit is determined by estimating a slope (*slope_vector*) from the difference of the averaged present energy level of the neighbours (*activation1(N1-4*)) and its own present activity (*activation1*). The *slope_vector* is than added to the *slope_old* of the processing unit under consideration to update the current state of the slope. The updated state of the slope is added to the *activation1* to generate the current energy level (*activation0*). This represents an integration process with 3 levels of processing of information. The first step is a spatial derivation is calculated with no temporal history. In the second step the value of spatial derivation is integrated by adding the derivation to the history of derivation (temporal integration). In the third step the updated history of slope is added to the history of energy level in order to determine the current energy state (second temporal integration). The derivation of *activation0* is a temporal derivation and results in a time dependent function. The second derivation of *activation0* is again a temporal derivation but results in a space-dependent function (all parameters shown in the microcircuit in **Fig. 2c**).

*Neighbour_integration (NI_slopev*) modulates the energy transfer of neighbouring neurons and in this way the extent of energy update by the *slope_vector. NI_slopev* operates like a low pass filter, whereas *damping* reduces the energy level, thus reducing propagation of oscillations. *Slopeo_damping* is not impairing propagation, but the duration of resonance oscillations of the system (normal mode or eigenschwingung). Besides those symmetrical calculations, modulation of *ratio_neighbour_activation* and *ratio_inhibition_activation1* enable alternating influence of the inhibitory dendritic subtraction and the excitatory influence of the neighbours (**Fig. 2c**). Increasing the GABAergic influence induced fast gamma waves (**Fig. S4** and ^62–65^), whereas increasing excitatory, uncorrelated or integratory influence leads to system collapse (**Fig. S4, S5 and** ^66–68^). *Marginextradamping* implements an extra damping for the margin to inhibit wave reflection at a sharp non-processing border.

Each step of processing can be related to its own biological meaning and modulated in many ways, such as integration steps are mediated by excitatory neurons and derivation is set up by a combination of excitatory and inhibitory neurons. Damping of *slope_old* or *activation0* is a damping of the cell bodies and thereby showing similarities to tonic inhibition^69^ (**Fig. 2c**). The *calculation of the slope_vector* is rather regulated by an excitatory-inhibitory balance mediated by axodendritic inhibition, potentially near synaptic, and/or indeed synaptic phasic inhibition^70–72^.

Schizophrenia simulation is implemented by introducing uncorrelated synaptic information transfer at the 3 different steps of information processing shown in Figure 2c. The transmission of the *slope_vector, slope_old*, or *activation0/1* can be deregulated randomly at various ratios, causing dissociation and beta band decline (see **Fig. 4g&h**). The Alzheimer model is set up by generating randomly no functional units called lesions in the model. The lesions can grow in size. The lesions can decrease or impair information distribution over the entire model which decreases information content (see **Fig. 4f**).

The main algorithm processing the information each time step for each unit can be described mathematically. Assuming 8 neighbours with activation levels (*activation1(N1-8*)), activation of unit in focus (*activation1/activation0), slope_old, slope_vector, activation3, activation4, neighbour_integration (NI_slopev), slopeo_damping, damping, ratio_neighbour_activation, ratio_inhibition_activation1 and marginextradamping*.

### Main algorithm

1. activation3 = (activation1(N1)+activation1(N2)+acttivation1(N3)+activation1(N4))/4
2. activation4 = (activation1(N5)+activation1(N6)+acttivation1(N7)+activation1(N8))/4
3. slope3 = activation3 – activation1
4. slope4 = activation4 – activation1
5. slope_vector = (slope3 + slope4)/2
6. slope_old = slope_old + slope_vector
7. activation0 = activation1 + slope_old
8. activation_change: within the main algorithm activation1 is the present energy level.

Modulation outside of the main loop is done on activation0. It is important to differ between those equal variables, as the processing units are queried several times within a processing step.

The main algorithm is resulting in **eq. 1**.

**Equation 1 (modulation see supplement):**

*i* = time steps
*k* = number of neighbours neighbours
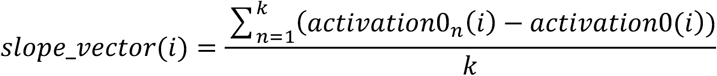
*slope_old*(*i* + 1) = *slope_old*(*i*) + *slope_vector*(*i*)
*activation*0(*i* + 1) = *activation*0(*i*) + *slope_old*(*i* + 1)
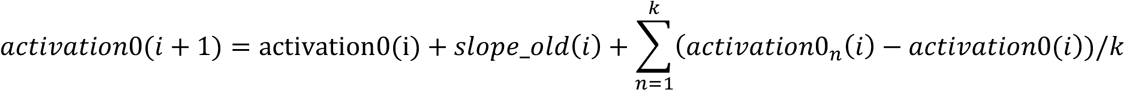

### Specific parameters of our simulations as shown in the figures

**Fig. 2b:** Continuous peak input frequencies are 13, 17, 23, 29, 31, 37, 41, 43, 47, 53, 59, 61 Hz. Further parameters are set as follows: *NI_slopev* = 1.2, *slopeo_damping* = 0.01, *damping* = 0.0001, *marginextradamping* = 2, *inputstrength* = 120 mV and simulation length = 3000 ms.

**Fig. 3a:** EEG signal: 21 sine-like stimuli of 400-500 Hz with 800mV amplitude and a random onset of bursts (average 250 ms) of 50 ms length are applied. 20% of stimuli were superimposed on a carrier of 0-30 Hz with 2400 mV strength. The input of slow wave state and waking state are alike despite an increase of *NI_slopev* from 0.1 to 2.6655 accompanied by a slight increase of *slopeo_damping* (1.0E-5➔1.0E-4) and *damping* (1.0E-3➔1.0E-2) decreases the amplitudes of waves and facilitates gamma oscillations. Raising the carrier frequency from 0-13 Hz to 0-23 Hz favours the usage of the resonance frequency (compare baseline in Fig. 4c and S3). In slow wave state low carrier frequencies enhance the effect of amplitude increase. The decrease of *NI_slopev* functions like a low pass filter. The EEG electrode radius is 10 mm.

**Fig. 3b&c:** Positive peak distribution of the MEA signals for slow wave state and waking state are indicated in Fig. 4b. Parameters are as described in Fig. 3a. Turning from waking state to anesthesia urges an increase of *slopeo_damping* (1.0E-2 ➔ 1.0E-1) and *damping* (1.0E-4 ➔ 1.0E-3). The *NI_slopev* is similar to waking.

**Fig. 3d**: Generation of overtones uses 7 Hz continuous peak input, a *NI_slopev* of 2.6655, *slopeo_damping* of 0.01, *damping* of 0.0001, *marginextradamping* of 2, *inputstrength* of 800 mV and a simulation length of 3000 ms. Measurements were done on LFP signals, simply generated by a small EEG electrode size (radius = 1 mm).

**Fig. 3e**: Wave speed estimate by 7 Hz continuous peak frequency. Overtones were generated which enabled the detection of maximum frequency that the simulation processed. The speed of the wave hills were altered in 7 steps (*NI_slopev* = 0.01, *NI_slopev* = 0.2, *NI_slopev* = 0.5, *NI_slopev* = 1.2, *NI_slopev* = 2.0, *NI_slopev* = 2.5 and *NI_slopev* = 2.6655) resulting in wave speeds of 0.03, 0.13, 0.22, 0.34, 0.45, 0.49 and 0.53 m/s. Other parameters were set as follows: *slopeo_damping* of 0.01, *damping* of 0.0001, *marginextradamping* of 2, *inputstrength* of 800 mV and a simulation length of 3000 ms.

**Fig. 4a**: Parameters for estimating a potential encoding >50 bit: 29 continuous peak prime inputs of 5-239 hz, *NI_slopev* = 2.6655, *slope_damping* = 0.01, *damping* = 0.0001, *marginextradamping* = 2, signal length = 3000 ms. Analysis was executed on MEA signals.

**Fig. 4b**: For performance estimate of model size 12 continuous peak prime stimuli were applied (13, 17, 23, 29, 31, 37, 41, 43, 47, 53, 59, 61 Hz). Matched frequencies, overtones and noise (artificial frequencies) were investigated. X-axis: model size in number of processing units (each 0.5 mm). Y-axis: matched input frequencies as number of detected overtones and noise. Further parameters: *NI_slopev* = 2.6655, *slopeo_damping* = 0.01. *damping* = 0.0001, *marginextradamping* = 2.0 and signal length = 3000 ms. Analysis was executed on MEA signals.

**Fig. 4c**: Baseline is generated applying solely spontaneous activity of 100 mV and 0.01% of activated processing units each time step. Further parameters are as follows: *NI_slopev* = 2.6655, *slopeo_damping* = 0.01. *damping* = 0.0001, *marginextradamping* = 2.0 and signal length = 10000 ms. Here, the EEG signals was analysized (diameter = 2 cm).

**Fig. 4d**: Spontaneous activity and coherence of the human brain mirrored. MEA coherence measurement of spontaneous activity (0.01% activated cells firing with 100 mV). The signal length is 10000 ms. The *NI_slopev* is 2.6655, *slopeo_damping* is 0.01, the *damping* is 0.0001 and *marginextradamping* is 2.0. The distance of the MEA electrodes is 0.5 (MEA1-2), 2 (MEA1-2), 4 (MEA1-3), 8 (MEA1-4), 16 (MEA1-5), 32 mm (MEA6).

**Fig. 4e**: Size distribution of positive peaks in the EEG signal (2 cm in diameter) is analyzed according to the number of neighbours building a directly connected processing units (*k value*). Parameters are the same as in Figure 4d, but the number of neighbours is altered as indicated in the figure legend.

**Fig. 4f&g**: The Alzheimer’s Disease and Schizophrenia simulation utilize the same starting parameter space, as described in Fig. 4b. In difference, in A.D. the x-axis shows the alternating ratio of lesion and in Schizophrenia the x-axis shows the variable ratio of uncorrelated processing. The analysis was executed on MEA signals.

**For beta band decline analysis in Schizophrenia** the input was of sinus wave type. Information is transmitted in bursts of, e.g., 50 ms. The information is superimposed on a carrier or reference signal and the information is coded in high frequency space (>400 HZ). The information and the reference beam can be complex (several different frequencies etc.). The signal is strong projecting on the spontaneous activity, a theta band and beta band will self-organize **(Figure 4h(left))**. Downsizing of the different derivation and integration steps (decorrelation) to random values between 1 and 0.8 in 99% of the columns regarding *uncorrelated4_slopeo, uncorrelated1*, and *uncorrelated_input*, as well as 12.5% of the columns referring *uncorrelated3_slopev* results in the beta band decline **(Figure 4h(middle))**. In **Figure 4h**(right) the delta is shown. Further parameters: *NI_slopev* = 2.6655, *slopeo_damping* = 0.01. *damping* = 0.0001, *marginextradamping* = 2.0 and signal length = 10000 ms. The EEG signals were analysed using a radius of 1 cm.

For generating **ripples,** parameters are as in the Schizophrenia simulation, and the carrier frequency is increased to 31 Hz (**Fig. S8**).

***In vitro* field recordings of murine hippocampal slices.** Data from^73^ are analyzed comparing carbachol induced gamma oscillations in wt mice and a constitutively active form of the erythropoietin receptor in GABAergic neurons **(VEPOR^+/+^)**^73^. Field oscillations were induced with 20 μM carbachol (Sigma-Aldrich) in transverse 400 μm hippocampal slices from male mice (27–33 day-old), protocols in^74^. Extracellular recordings were performed in the *stratum radiatum* of CA3 at 33°C in a Haas-Top interface chamber, using a MultiClamp 700B amplifier. Data were sampled at 10 kHz using an Axon Instruments Digitizer 1440A with pClamp 10 software (Molecular Devices, Sunnyvale, CA, USA). The analysis was equivalent to **Fig. 3b,c** for baseline activity prior to carbachol application and for 0-15, 15-30, 30-45, and 45-60 min time segments after carbachol application. Results are displayed in **Fig S9a (wt)** and **S9b (VEPOR^+/+^)**^73^.

Parameters as in **Fig. 4h** and **Fig. S8** demonstrate **the effect of undersampling (Fig. S10)**. The stimulus onset was random. 21 different stimuli per second using an information bandwidth of >400 Hz were superimposed on carrier waves ~5 Hz. The location of onset was also random. The signal was smoothed in frequency space with a running average using a window size of 120 datapoints. The signal length was 8000 ms.

