## supplement for "Quantifying and modelling non-local information processing of associative brain regions"

**Supplementary Material:**

**Information is processed on ever higher levels and networks.**

(i) We give first the details of the **DNA master equation** (**level 1; pp. 1-2**).

(ii) Our DNA encodes all protein networks of the body, a representative signalling network simulation for neurons is shown, the **BDNF/TrkB signalling cascade** (**level 2; pp. 2-6**).

(iii) Encoded by the neuronal protein network, differentiated and active neurons emerge from this, leading to non-local wave-like patterns. Details of our **brain simulation** (**level 3; p. 6-25**) encapsulating these and their emergent information integration properties are shown and how well the simulation correlates with observations from neurobiology and anatomy. For installation and program details see our Tutorial provided as additional file.

(iv) We close this file with a critical appreciation of **network biology** (p. 25) as a possible "theory of everything" and inherent limitations.

**Level 1: DNA encodes all networks:** (i) Storage of internal information in the form of genes and gene regulation as well as all higher levels of information storage are central to *life*: A master equation (**eq. S1**) is introduced to calculate increase and decrease of different DNA sequences representing a quasi-species in viridae or higher levels of information storage (genomes or a community of several genomes) in more complex organisms. (ii) Higher levels of information storage do no longer just rely on the DNA sequences but involve subsequent transcription into RNA and translation into proteins. Even higher levels store information in protein-protein networks, for instance differentiation of a cell into different cell types. Multicellular organisms develop different organ-specific cell types including neurons. Neuronal networks operate then on a higher level between neurons and allow besides genetic information the storage of experiences or learned behaviour. Only with this higher neuronal level, an organism-specific "*meaning*" arises in the form of adaptive behaviour to satisfy drives and on a still higher level understanding of the environment, for instance successful hunting behaviour or planning an attack. Higher levels of components and interactions always imply higher-level modules, such as neuronal cells as a new level, provided that genes are there to encode different types of neurons. This new higher level of modules also allows new behaviour of these modules, in the example the new learning behaviour of a neuronal network. Network biology can in this way define "*emergence*", the arising of qualitative new properties by examining ever higher-level modules as new types of sub-networks. (iii) Network biology is hence applied here in different examples to study both simple and rather complex organisms, as well as environmental and host-pathogen interactions, looking at their regulatory and metabolic interaction networks up to different species interactions, ecosystems including human civilisation. To derive the master equation, we combine two basic principles for living beings: First, information for survival is selected by evolution. Second, this information is stored internally on higher and higher emergent levels: Basic is hence **the master equation** on DNA and its evolution governed by replication (r), mutation (m) and selection (s):

$$\frac{d}{dt} * DNA_{vector}i = DNA_{vector}i * \left(r * p(r)\right) * \left(1 - p(m)\right) * (1 - p(s))$$

(**eq. S1**)

Encoded in DNA, there are next molecular networks. On the next level this leads to inter- and intracellular communication, different cellular phenotypes and cell types and neuronal network behaviour (depicted in **Fig.1a-e)**.

Individuals come together and form the human society. Higher levels of emergence are thus described by equations modelling metabolic and regulatory networks (e.g. [1,2]), cellular circuits (see next chapter), brain simulations (below) and then organismic interactions in ecosystems or even in a human civilisation. Each higher level develops new properties not evident from the lower level componentes, in particular regarding information storage and processing as well as functional capabilities to do so, i.e. to analyze and respond to the environment.

**Level 2: Gene-coded signalling network**

Our model describes the emergence of higher neuronal functions from DNA encoded wiring of neuronal circuits resulting in non-local processing of information. On level 2 cellular concepts of signalling networks are introduced (**Fig. 1 a-e**). This is exemplarily shown for the BDNF directed signalling cascades and how this supports multiple cellular functions (**Fig. 1b, c, e; Fig. 1**) and how it even causes differentiation of subtype-specific neural cells. The cellular simulation allows easy and explicit testing and incorporation of the effect of different mutations on the DNA or protein level. It is also suited for simulations regarding the transcriptome or protein-protein interactions including systems effects implying differentiation into different neuronal types or types of synapses. This includes general impairments of cellular network function implying defects of higher network activity and disease (e.g. Alzheimer´s and Schizophrenia).

The BDNF/TrkB signalling network was iteratively modified to account not only for the direct activation route of the TrkB receptor by its natural high-affinity ligand BDNF[3], but also for different neurotrophin-independent TrkB transactivation routes[4,5] (**Fig. 1b, c**). **All nodes, interactions, references** (see excel **Table S1**): Gene-coded protein networks and their intra- and intercellular signalling functions influence the response behaviour of neural cells. Even simple forms of activation of neural cells (neurons or glial cells) reveal forms of wave-like information processing (depicted in **Fig. 1d**). Validation data include induced neuronal network activity in cultured neurons (Video 1; available as supplement and at https://www.biozentrum.uni-wuerzburg.de/bioinfo/computing/neuro; example taken from[6]). In this example, hippocampal neurons underwent a chemical-LTP (long-term potentiation) stimulation. Chemical activation of neurons induced synaptic communication between neurons and developed synchronized activity over all neurons in the field of view. Visualization of the phenomenon bases on changes in intracellular calcium concentration that can be seen in numerous loci over neurons (**Fig. 1d**, white circles indicate loci of calcium activity that can be computed and evaluated with modified wavelet algorithms, as described recently[6]).

Other simple forms of wave patterns of cellular signals taking place in biological models are observed in glial cells. Glial cells generate signals in waveform, but the cells and signals are functionally different from those observed in neurons. In glial cells, the neurotransmitter glutamate induces calcium waves, a phenomenon of long-range glial signaling[7]. This is exemplarily shown in Video 2 (available as supplement and at https://www.biozentrum.uni-wuerzburg.de/bioinfo/computing/neuro. Here, glial cells cultured from a postnatal day 4 mouse were loaded with a calcium indicator and calcium signals were imaged with a video camera, as described earlier[6]. Cells were stimulated for a short time (900 ms) with 500 µM glutamate to activate metabotropic glutamate receptors. Cells were recorded at 10 Hz. Local application of the glutamate was performed with a patch pipette. In this video for glial cells, the wave dispersion pattern of the cellular activity can be clearly observed. These signaling patterns are also observed in brain tissue[8]. Glia calcium waves are per se activity waves[9], but are different from neuronal oscillations (Theta-, gamma)[10].

Our work shows that a disruption in such patterns is a symptom and/or a cause of several pathologies in the brain (Fig. 3 and 4 in results). The wave like pattern activity of the brain occurs both in space and in time. Similar spatial and temporal dependencies are known from the activity bands in electroencephalographic signals.

In summary, Fig. 1 illustrates how wave-like processing patterns result from molecular wiring for neural cells. Simulation of inter- and intracellular protein networks show how neural cells develop into different directions in dependence of signalling events, as indicated by the simulation of neurotrophin-independent TrkB activation effects. The videos at https://www.biozentrum.uni-wuerzburg.de/bioinfo/computing/neuro show in detail how neural cells react on activity stimuli. This has then as a new emergent effect a) synchronous activity (Video 1) and b) coupled activity with resulting wave-like patterns of activity (Video 2).

These observations motivated us to translate both features into the neurobiological simulation (see eq. 1 in the results) and then to investigate which higher-level emergent effects could come from this synchronous activity and/or the coupled activity of many neurons (see level 3, neurobiological model).

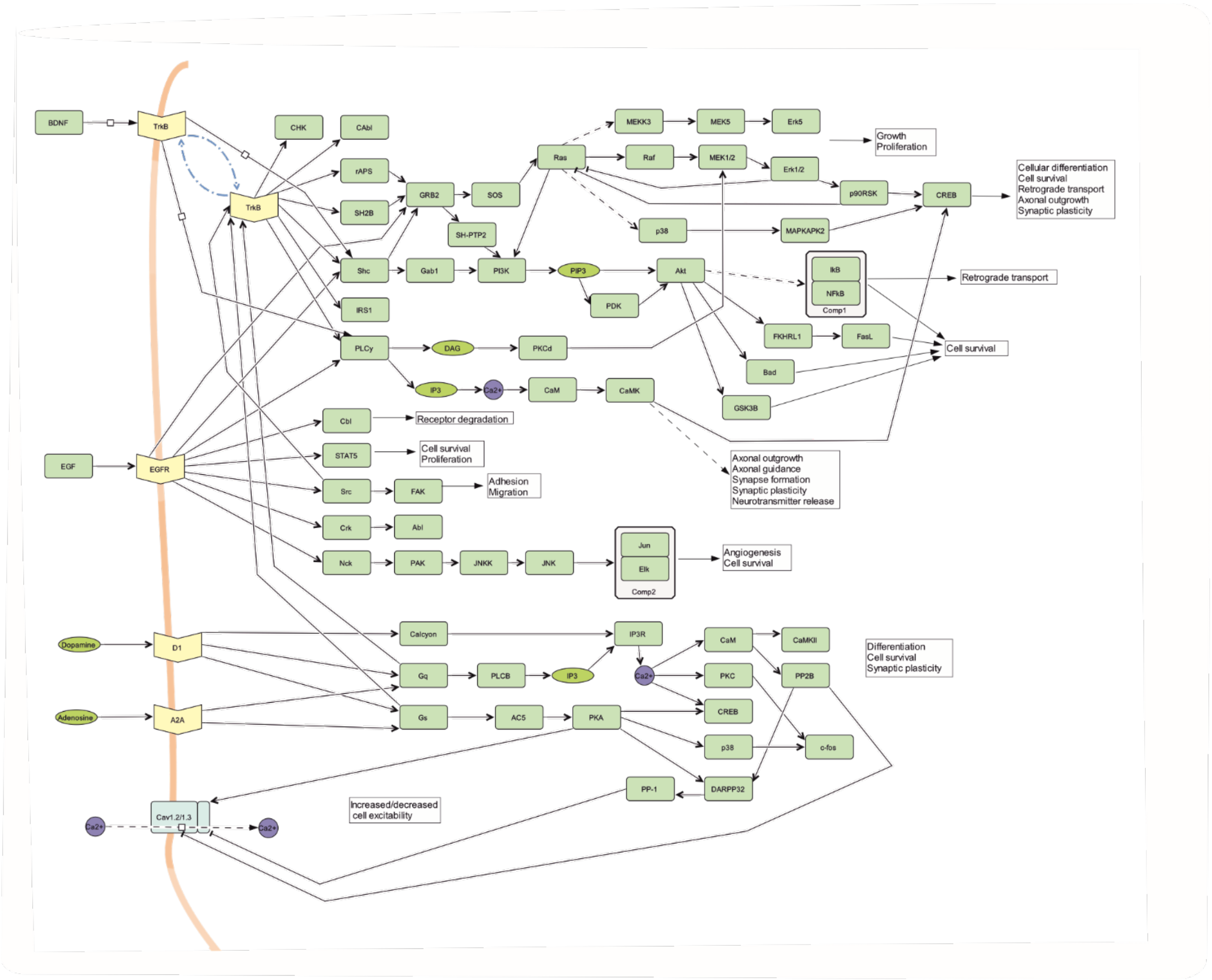


**Fig. S1: (A) Cellular model of the BDNF / TrkB signalling network:** This model (protein network, level 2) is presented in **Fig. 1b, c** and is given here in detail. The proteins for this are encoded on the DNA level (basic level 1). It is based on Table S1 (all nodes, interactions, references; hand curated) and was assembled starting from the KEGG pathway "ko04722 Neurotrophin signaling pathway"[11,12]. This pathway presents the direct activation mechanisms of the neurotrophins NGF, BDNF, NT3 and NT4 and the subsequent activation of the intracellular signalling cascades including the pathways represented by MAPK, PI-3 kinase and PLC. The activation of these cascades results in functions like axonal growth, cell migration, cell survival, cellular differentiation and plasticity. The pathway was later modified to allow for different neurotrophin-independent TrkB transactivation routes. A first transactivation route starts with charged ions like zinc or calcium[13] passing through channels like VDCC or NMDAR. Another starting factor of TrkB transactivation is dopamine[14], which activates G-protein-coupled receptors and subsequently transactivates TrkB receptors. A third TrkB transactivation route is mediated by adenosine and the A2A adenosine receptor, another G-protein-coupled receptor[15]. Finally, the growth factor EGF is also present as a TrkB transactivation factor[16].

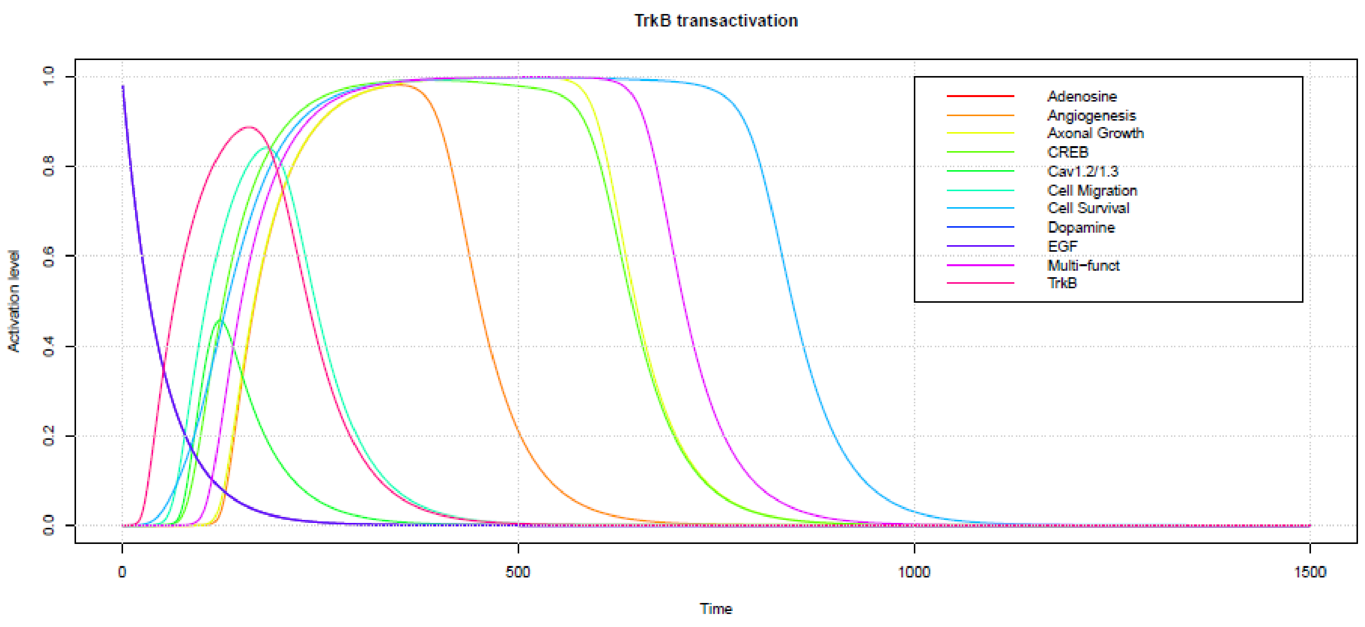


**Fig. S1(B), panel 1**

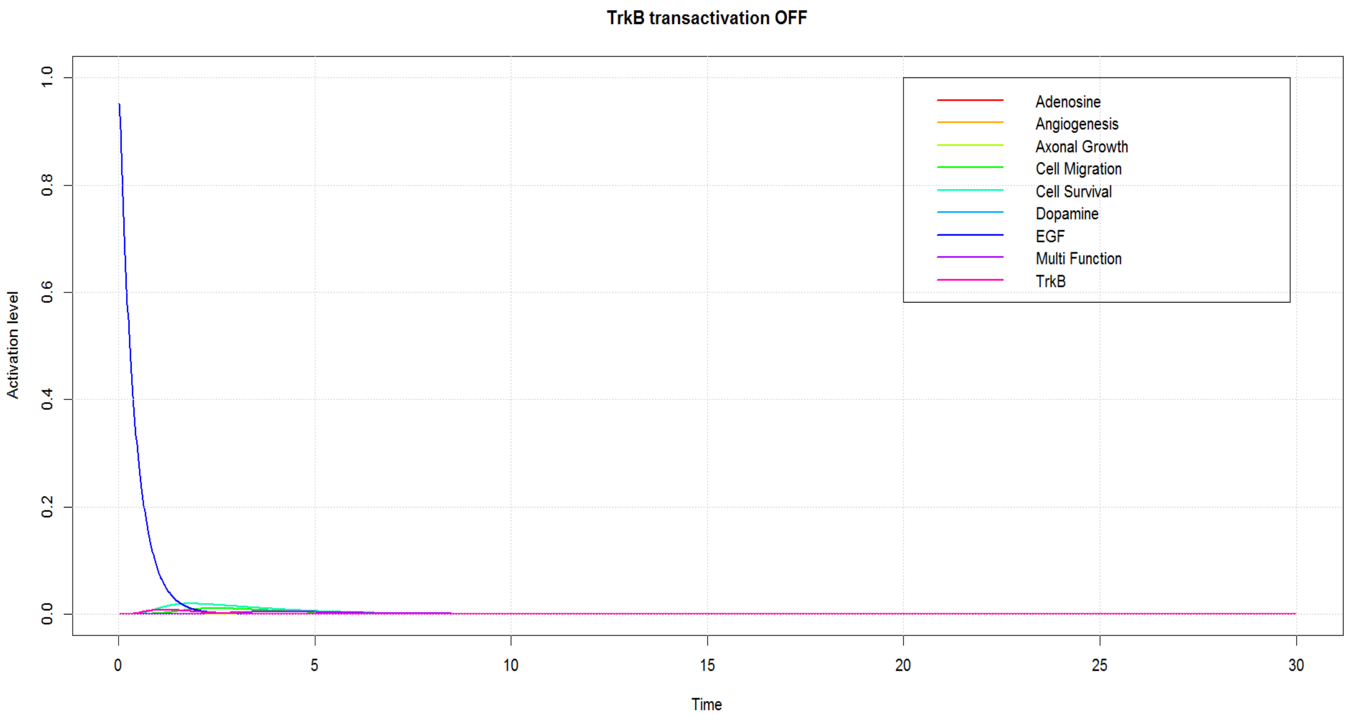


**Fig. S1(B), panel 2**

**Fig. S1 (B) Cellular phenotypes including differentiation arise from modulating protein-protein interactions between cell compartments.** Extended version of **Fig. 1e**. The simulation illustrates how BDNF signalling versus neurotrophin-independent TrkB transactivation can result in different cellular responses and thereby affect multi-functional signalling cascades, but also functions such as axonal growth or cell survival. Important key factors such as the activity-dependent transcription factors CREB or c-FOS are pointed out. The simulation shows different results depending on whether neurotrophin-independent TrkB transactivation routes induced by the adenosine receptor A2A (A2A-R), EGF-receptor, or dopamine receptor D1-R are ON (panel 1) or OFF (panel 2). Note that neurotrophin-independent TrkB activation can occur at intracellular sites and defined cellular functions no longer emerge when the decay factor of TrkB activation is changed. The neuronal protein-protein network simulation was run using the topology of the BDNF signalling network shown in **Fig. S1a** and its dynamic behaviour is in line with experimental data. For instance, A2A-R-dependent activation of TrkB can contribute to neuronal survival, at least in motoneurons and blocking this transactivation route can reduce the A2A-R mediated effects [15].

### Level 3: Neurobiological Model - Holographic non-local circuit (Netlogo simulation)

Originating from DNA storage and subsequent generation of neuronal circuits, the emergent level three results in cortical organization of neurons and wave patterns measurable by EEG. These are non-local wave patterns with holographic properties. This is described by our high-level simulation and explained now. We show here (i) how the modulation of this circuit and its emergent properties can be directly achieved by looking at different neighbour interactions and the resulting changes in the wave patterns. (ii) We next analysed the electrode signal in the extracellular space around neurons and show again that the simulation agrees well with observation and experimental EEG data. To confirm this, we look at basal brain activity, beta firing and inhibitory neurons, epilepsy and non-local information and include an analysis of the peak half-width. Finally, the coherence of the stimuli is investigated. This again shows strong parallels between recorded EEG data and the analogous phenomena produced by our holographic simulation.

#### (i) Modulation of the neuronal circuit

For the following it is useful to look at the components of **eq. 1** given in the results. It has to be emphasized that the influence of neighbours on the *slope_vector* in the simulation is typically 1 to 8 (simplification: direct neighbors on a grid). This symmetrical behaviour can be changed by the influence of neighbour energy (*ratio_neighbour_activation*) and the subtraction of the own energy (*ratio_inhibition_activation1*). The contribution of the calculated *slope_vector* on the update of *slope_old* is modulated by *NI_slopev*, meaning that activity of 8 neighbours and its derivation from the own activity can be summed and transmitted to the central neurons, or normalized by the extent of *NI_slopev*. Likewise, the contribution of *slope_old* to *activation0* can be modulated. We assume here a symmetrical calculation of the neighbour energies and the following calculation steps but suggest to consider asynchronous and individual modulation of each kind of neuronal associations. The modulation is also displayed in a neuronal microcircuit (see **Fig. 2c**).

1. slope3 = (activation3 * ratio_neighbour_activation – activation1 * ratio_inhibition_activation1) * neighbourintegration;

2. slope4 = (activation4 * ratio_neighbour_activation – activation1 * ratio_inhibition_activation1) * neighbourintegration1;

3. slope_vector = (slope3 + slope4) / nc

4. slope_vector = slope_vector * NI_slopev

5. slope_old = slope_old / (1 + slopeo_damping)

6. activation0 = activation / (1 + damping)

7. activation0 = activation0 / (1 + damping + (marginextradamping / (1+ minimum_margin)))

Marginextradamping increases with distance of margin to the centre of the model.

The performance, the coding, and execution of the simulation and these steps were tested independently on a computer cluster by expert computer scientists, who in addition to this, established a parallelized version of the simulation for the cluster.

By these rules, the simulation then creates wave patterns that arise on a high level. The wave patterns connect all Mountcastle columns in a large area, providing the same information everywhere. Program code deposited at https://www.biozentrum.uni-wuerzburg.de/bioinfo/computing/neuro .

The *in silico* modulation of the processing steps 1-7 above reproduces typical EEG phenomena, such as slow waves in sleep and EEG patterns of the awake state. Results in **Fig. 3** and **Fig. 4** show this also for different pathophysiological conditions (e.g. in Schizophrenia, Epilepsy, Alzheimer's disease, anaesthesia). In the following supplemental **Fig. S2** till **Fig. S10** and their legends provide more details on the simulation and comparisons to pathophysiological conditions and these results are discussed (p.16-21) including parameter values, coding capacities and direct comparison with experimental data from literature.

The resulting modulated main algorithm (modulation, steps 9-15) is given in **eq. S2**.

**Equation S2:**

$$slope\_vector(i) = \frac{\sum_{n=1}^{k}(activation0_n(i) * ratio\_neighbour\_activation_n(i) - activation0(i) * ratio\_inhibition\_activation1(i))}{k} * NI\_slopev$$

$$slope\_old(i+1) = (slope\_old(i) + \mathrm{slope\_vector(i)})/(1 + slopeo\_damping)$$

$$activation0(i+1) = (activation0(i) + slope\_old(i+1))/(1 + damping)$$

$$activation0(i+1) = \frac{\text{activation0(i)} + \dfrac{slope_{old(i)} + \sum_{n=1}^{k} \dfrac{activation0_n(i) * ratio_{neighbour_{activation_n}}(i) - activation0(i) * ratio_{inhibition_{activation1(i)}}}{k} * NI_{slopev}}{1 + slopeo_{damping}}}{1 + damping}$$

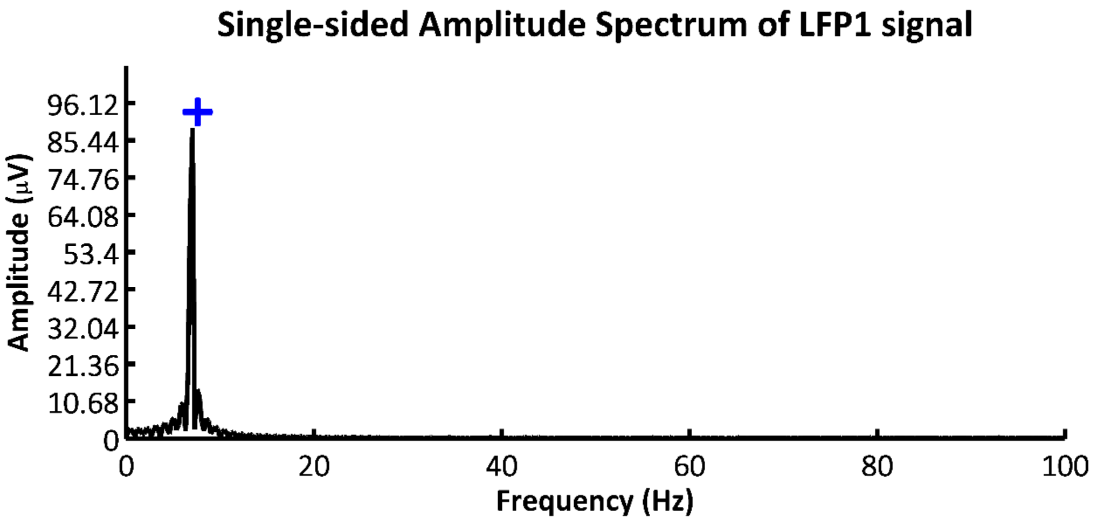


**Figure S2: Sinus rhythm extinguishes LFP harmonics in line with experimental observation.** Shown is the analysis of overtone decline in presence of sine input generated by the simulation. The cortical architecture of the brain (rodent hippocampus) can produce non-local overtones from short pulsed peak input, sinusoid input decreases them (compare to Laxpati *et al.* 2014[17], Figure 7). The comparison of **Fig. 3d** and **Fig. S2** indicates the overtones decline using sine input in silico that is in good accordance with experimental data[17]. For demonstrating overtone decline 7 Hz continuous sine input are applied, as well as a *NI_slopev* of 2.6655, *slopeo_damping* of 0.01, *damping* of 0.0001, *marginextradamping* of 2, *inputstrength* of 800 mV and a simulation length of 3000 ms. The LFP (local field potential) signal is defined as the summed activity of columns within a radius of 1 mm.

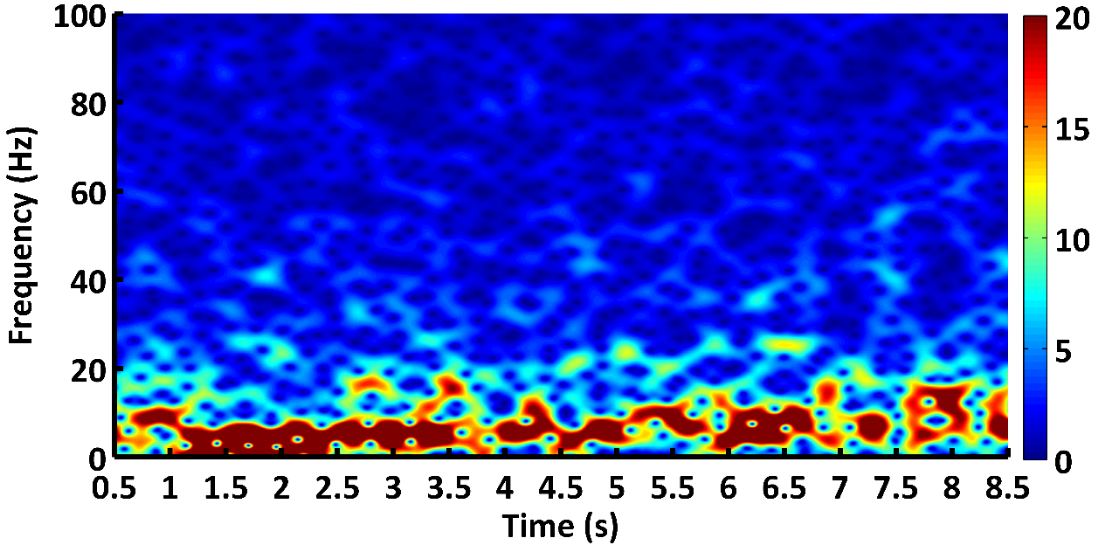


**Figure S3: Arousal: Basal brain EEG activity changed from theta to alpha activity.** For this, the Baseline change is mediated in our model changing the parameter *NI_slopev* (theta to alpha). The *NI_slopev* is stepwise increased from 0.1 to 2.8 by 0.3 each 1000 ms. The other parameters are the same as in **Fig. 4c**. The signal length is 10000 ms. Here, only the period from 0.5 to 8.5 s is shown. The EEG electrode radius is 10 mm. The transition from theta to dominating alpha activity in accordance with transition from sleep to waking is well observed in EEG recordings in men[18].

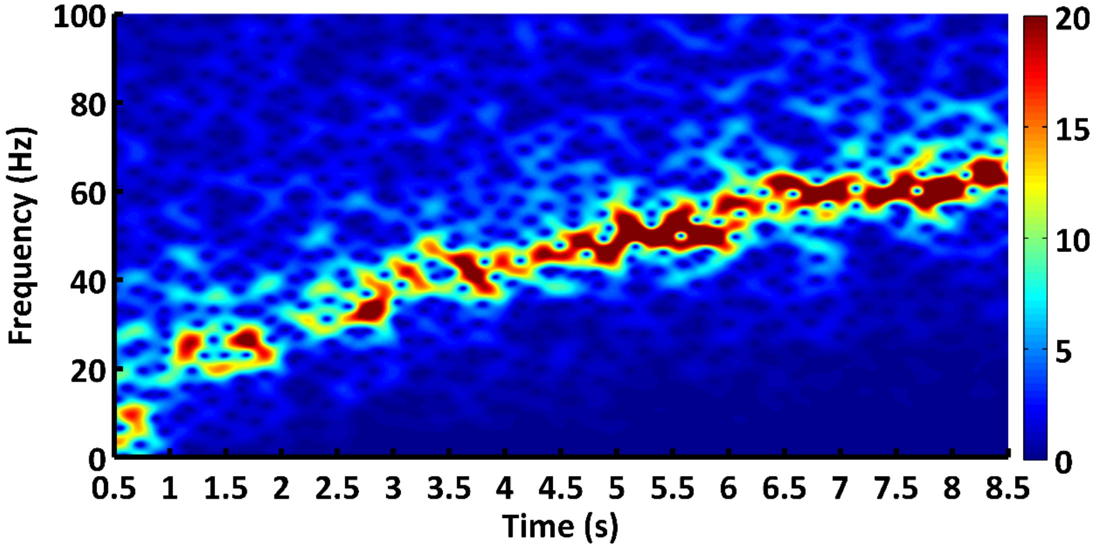


**Figure S4: EEG beta firing increases with increased relative activity of inhibitory neurons** (increase of *ratio_inhibition_activation1*). Using the parameter set up described in **Fig. S3 starting with an** *NI_slopev* of 2.6655, as well as a *ratio_neighbour_activation* (excitatory neurons) and a *ratio_inhibition_activation1* (inhibitory neurons) of 0.8, the *ratio_neighbour_activation1* is increased each 1000 ms by 0.02. In effect, the beta firing is increasing gradually which is also valid for cortical circuits[19-22]. Here, we used a simulation setup below criticality, as a similar set up would cause system collapse when going into super criticality, as described in the literature[23-25]. This shows that beta firing increases with more inhibitory neuronal activity in our simulation just by the emergent wave pattern effects in full accordance with experimental data[19-22].

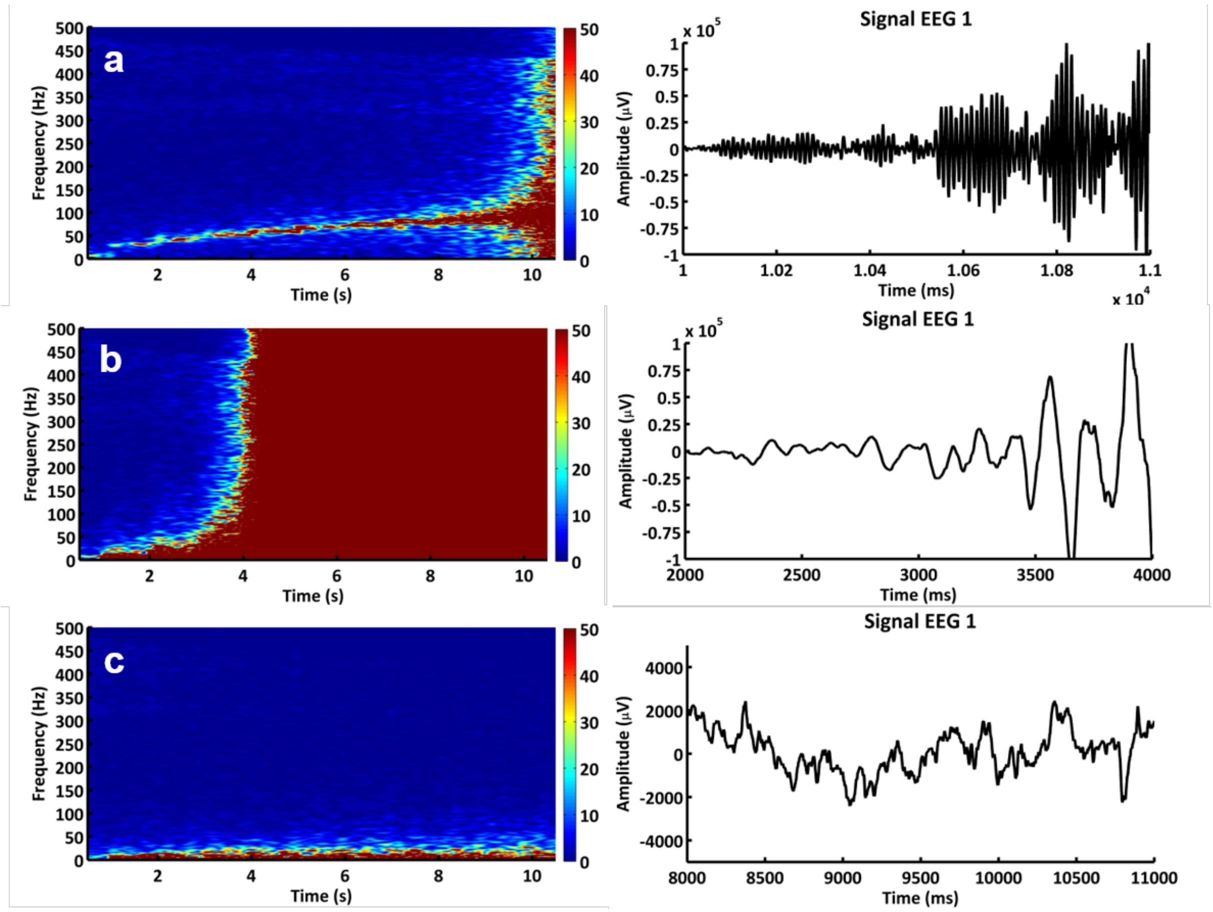


**Figure S5: Analyzing Epilepsy considering non-local information: Sharp border from subcriticality to supracriticcality[23,24].** There are several effects causing a system to collapse and produce infinite numbers, which is linked to a stepping from subcritical state to a supracritical state and subsequent linked to epilepsy[23]. Typically, the system state turning from subcriticality to supracriticality is defined by a sharp border[23,24]. For the non-local model we can show that a small increase of a *NI_slopev* that is close to criticality (2.6655) to a value of 2.7 causes the system to produce infinite positive and negative number. Further, the *ratio_neighbour_activation* shows a small tolerance to an increase to a level a little bit higher than *ratio_inhibition_activation1*. Even distant from criticality (*NI_slopev* = 2) a *ratio_neighbour_activation* value of 1.001 compared to *ratio_inhibition_activation1* of 1 causes the system to produce either only positive or negative infinite numbers (exponential function). Additionally, from **Fig. S4** we learn that the increase of *ratio_inhibition_activation1* is well tolerated. However, closer to criticality the likelihood increases that an increase of *ratio_inhibition_activation1* will also cause system collapse. Those results are not shown due to their clearness. Here in **Fig. S5** we describe a statistically uncorrelated change of the values *ratio_neighbour_activation*, as well as *slope_vector* and *slope_old*. The uncorrelated function is similar to that used for Schizophrenia simulation, which might also explain why Schizophrenic patients have an increased likelihood for seizure[26,27]. In **(a)** the *ratio_transmitted_energy* of the neighbouring neurons is gradually declined by 0.05 each 1000 ms and results in a value of 0.5. Uncorrelated means that e.g. a random value between 1 and 0.5 is estimated that determines the ratio of energy transfer of 50% of the columns. This is analogous to an increased influence of inhibitory neurons. In **(b)** the *slope_vector* and in **(c)** the *slope_old* is statistically declined likewise. An uncorrelated change of the extent of *slope_vector* and *slope_old* is not disrupting the balance of excitatory and inhibitory neurons, but the balance of integration steps. This can also lead to epileptoform waves **(b,c)** which might also explain the variety of epileptiform waves. Further, we learn that epileptic seizure is possibly associated with statistical uncorrelated energy transmission as shown in **(a),** as a decrease of the *ratio_neighbour_activation* alone is not producing infinite numbers[23]. In other words, a decrease of the excitatory/inhibitory ratio is not always producing epileptiform waves, but if uncorrelated to a distinct extent it does. This might align with pathology and causes of epilepsy, such as the neuronal disfunction in diabetes[28,29], Schizophrenia[26,27], or A.D.[30]. All signals are EEG signals with a radius of 10 mm.

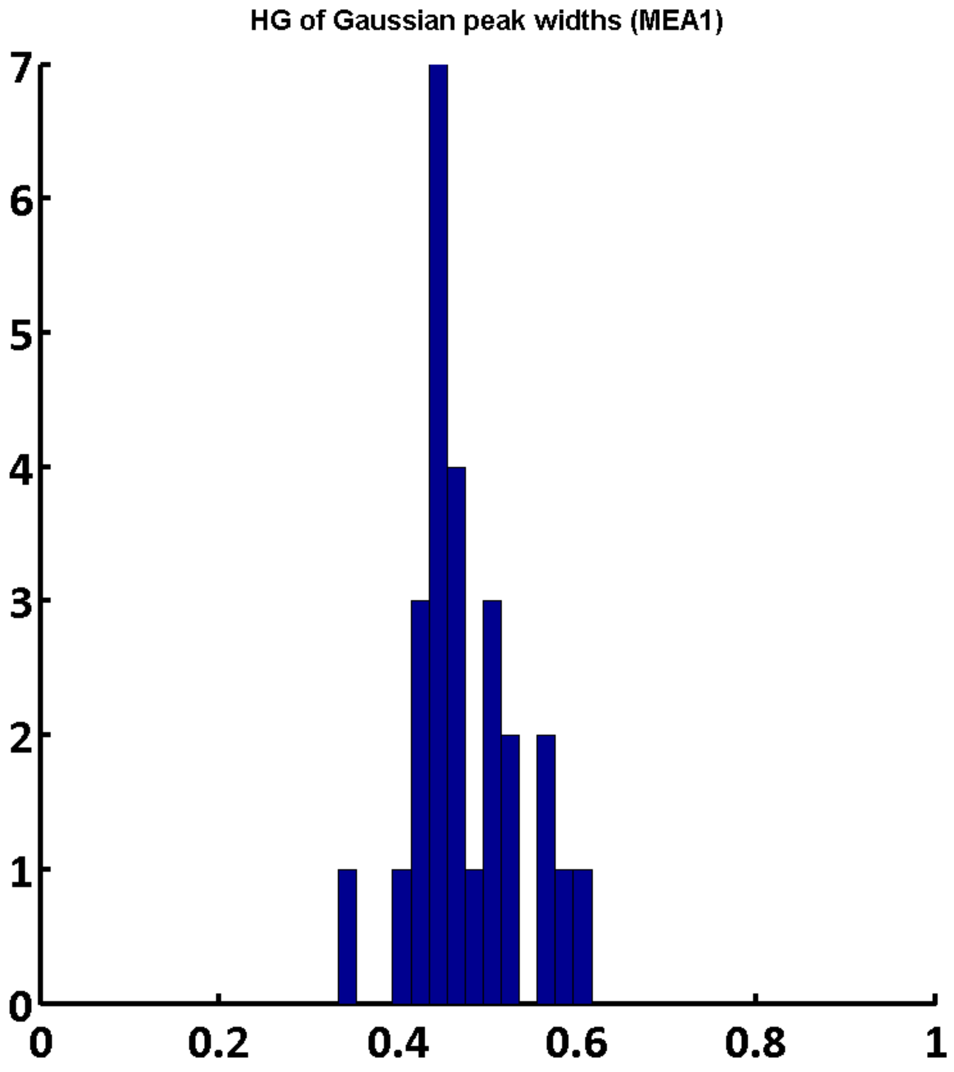


**Figure S6: Half width of the peaks.** Resulting from the peak analysis of the simulation of prime number frequency input from 2-239 Hz the half-width of the peaks is demonstrated. The input frequencies are continuously applied. The length of the analysed signal is 3000 ms. Other parameters used are the same as in the control of the Schizophrenia simulation (**Fig. 4h (left)**). The peak analysis shows that non-local information storage is efficient and works over the whole band-with. Using the equally distributed half-width of the peaks the coding potential within a distinct time frame can be estimated (200 bit within 1 second within a bandwidth of 10-110 Hz). The sharpness of the peaks improves by dividing the frequency space through its own running average (window size = 60 data points).

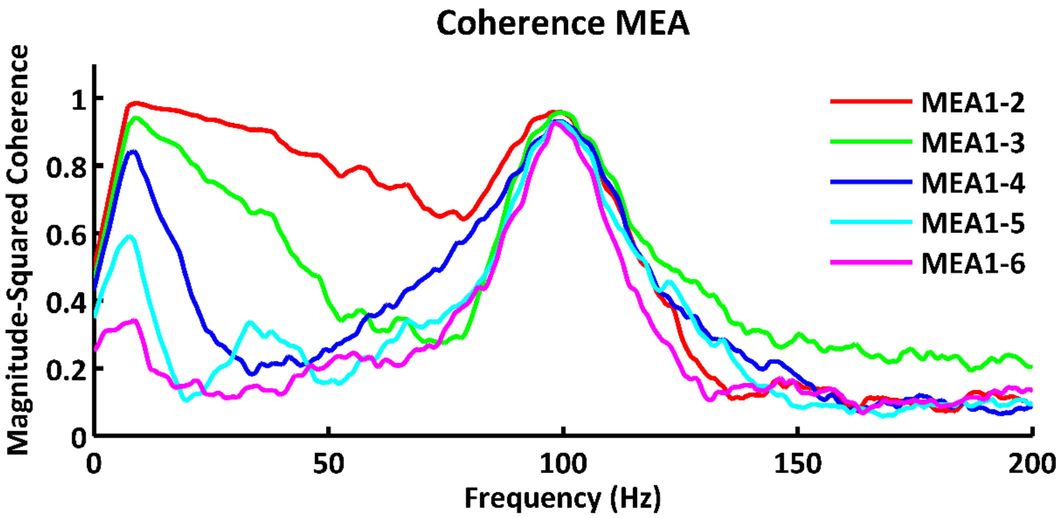


**Figure S7: Coherence and stimuli.** A sine stimulus of 100 Hz is set 500 ms after start of the simulation. The stimulus lasted for 200 ms. The coherence increases at the input frequency. Parameters are as in **Fig. 4d.** Coherence and stimuli response correlate well with the experimental data[31] .

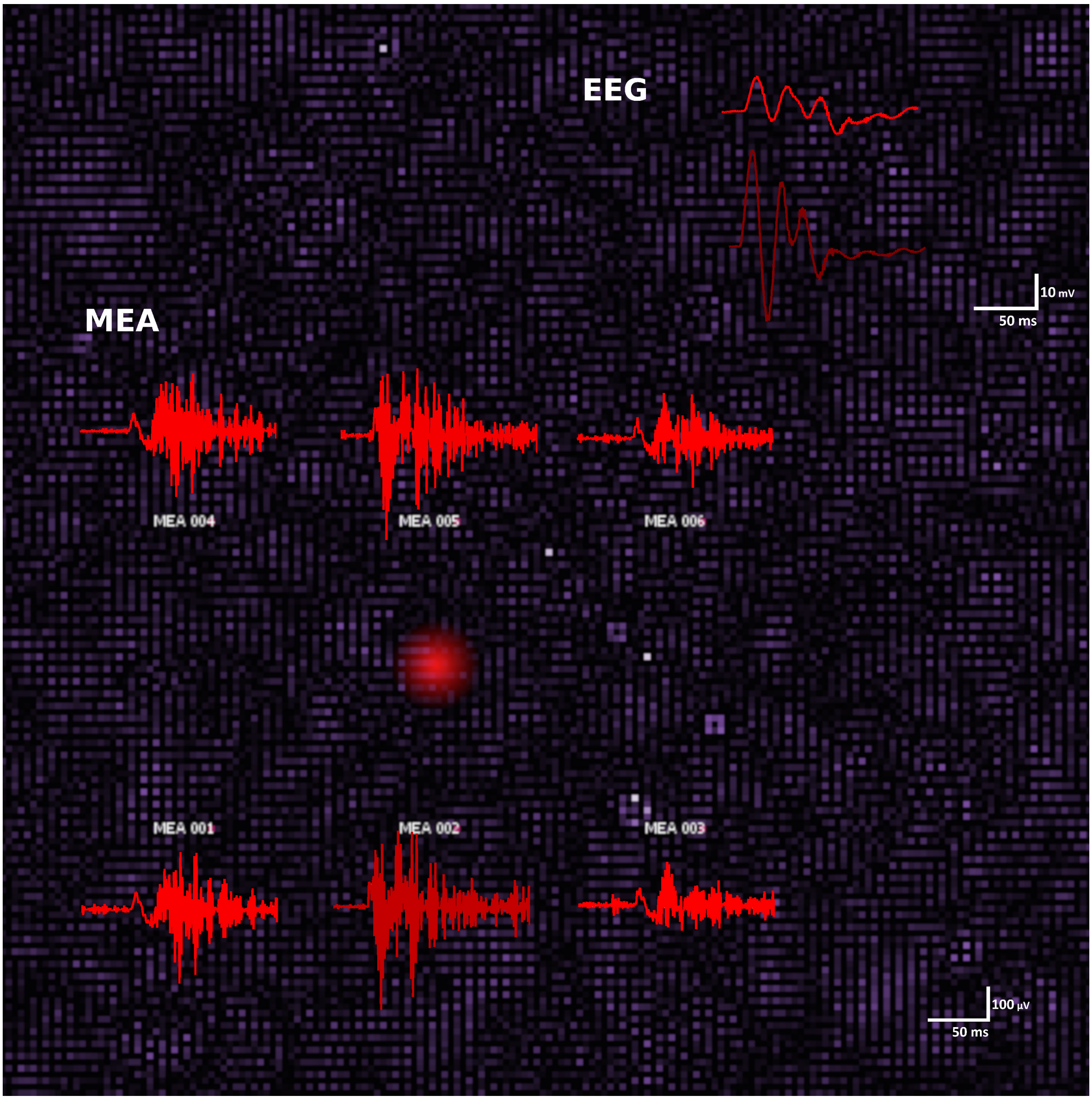


**Figure S8: Ripples in MEA signal.** We show here the activity of a simulated associative cortex region 500 s after stimulus onset (region of stimulus input indicated by red dot). The six MEA electrodes are indicated and the associated activity pattern during stimulation (stimulation period 50 ms). The related EEG signal is also shown. The paramters for the simulation are the same as described for **Fig. 4h**, except that the carrier wave has a frequency of 31 Hz.

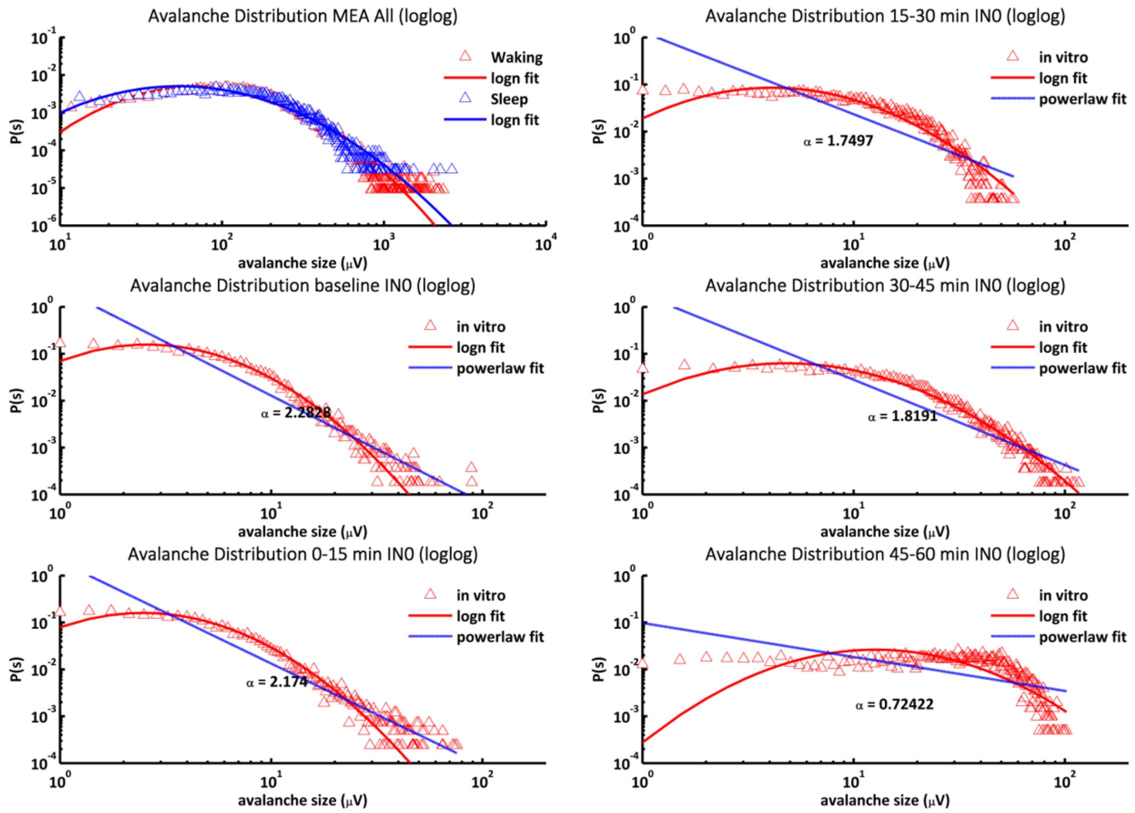


**Figure S9a: Comparing *in silico* and *in vitro* distributions.** Avalanche distribution of Fig. 3b in silico waking and sleeping state is indicating for direct comparison with *in vitro* recordings. We compare here electrode recordings of hippocampal brains slice of a 33 day old **wt** mouse. Shown are the baseline activity for 15 min and periods after stimulation with CCH (carbachol) of 0-15, 15-30. 30-45 and 45-60 min.

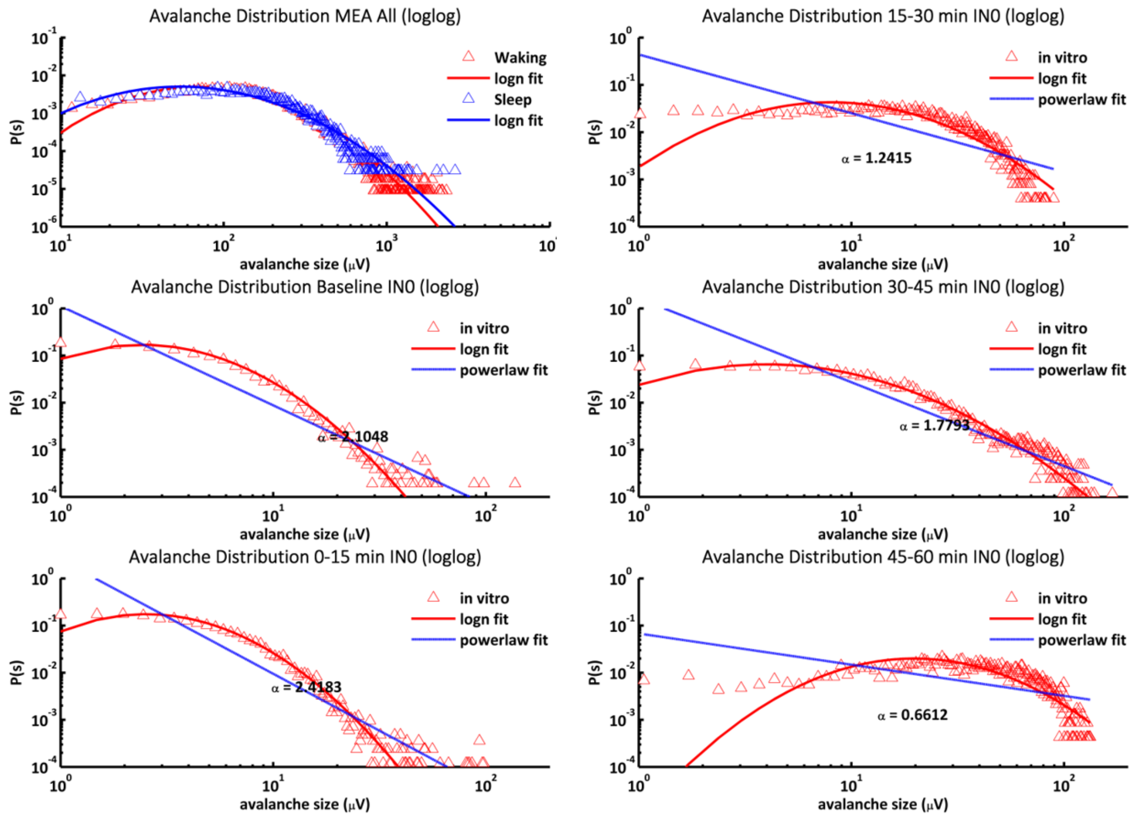


**Figure S9b: Comparing *in silico* and *in vitro* distributions.** Avalanche distribution of Fig. 3b in silico waking and sleeping state is indicating for direct comparison with in vitro recordings. We compare here electrode recordings of hippocampal brains slice of a 27 day old **VEPOR**$^{+/+}$ mouse[80]. Shown are the baseline activity for 15 min and periods after stimulation with CCH (carbachol) of 0-15, 15-30. 30-45 and 45-60 min.

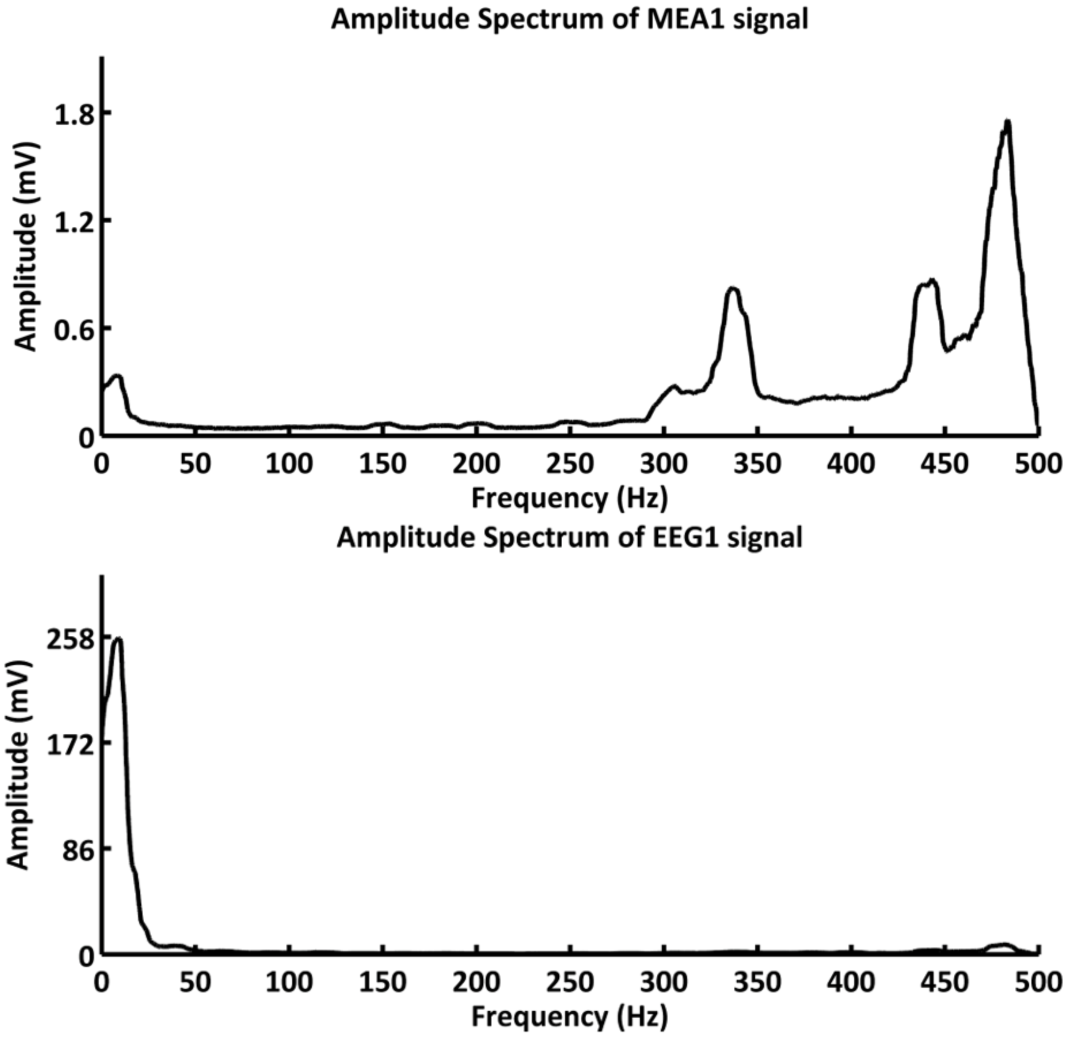


**Figure S10: Undersampling: suggests higher coding bands.** We show here that the decoding of a signal of an information bandwidth >400 Hz superimposed on carrier waves shows different distributions on small scale (MEA recordings) and on integrated scale (EEG recordings). Parameters are the same as describe for **Fig. 4h** and **Fig. S8**. Instead of 1 signal, 21 different signal per second were randomly applied. EEG suggests low frequency coding, whereas MEA point to high frequency coding. The effect is called undersampling.

**Further Discussion of brain features and their simulation in our non-local brain activity model**

**Emergent avalanche distributions** are very close to critical distributions[23,32-34,] to achieve maximum information content[35](**Fig. 4e**). The avalanche distribution of summed magnetic fields of columns (fMRI signal) is not only producing near critical distributions, but also shows an extended shift to higher activity by elevated connectivity[33]. Here, the summed activity of the columns within the radius of the EEG reproduces both.

**Key theories on consciousness**[36] are a long and complex subject. Here some central theories and authors are briefly presented for comparison to our approach and its explanatory value. It is important to include the global workspace theory[37] in which working memory is central. The center stage is illuminated by the focus of our attention, the consciousness, the remaining major memory is unconscious. Integrated Information Theory stresses Phi, the total information processing capacity of the brain as central. Given sufficient complexity this gradually leads to consciousness[38]. Eliminative materialists[39] see nothing special about consciousness, all is just neuronal activity. Illusionists focus on our highly parallelized brain[40] and denounce consciousness as a post-processing artefact of proprioception of specific actions.

**Further discussion of the simulations described in Fig. 3 and Fig. 4:**

The discrimination of the brain states **(results, Figure 3a-c)** reduces the complexity of the cortex to distinct energy coupling parameters related to criticality, emergence and non-locality. The anaesthesia state in the simulation is representing extrasynaptic tonic inhibition of the cell body combined with minor phasic inhibition affecting excitatory and inhibitory neurons likeswise[41-44]. In anaesthesia the first and second integration step within a processing unit (*slope_old* and *activation0*) are dampened. In other words, the history of slope and activation level are dampened via cell body damping. In effect, information is localized in anaesthesia causing a dissociation state. By contrast, although sleep also indicates inhibition, here the energy of information transfer between the neighbours is reduced (*NI_slopev*), while cell body damping is slightly reduced. Reduction of *NI_slopev* is an axodendritic damping and by this differs as it represents a current energy delta without historical predecessor. Sleep rather undergoes balanced inhibition of distinct excitatory and inhibitory synapses, axons, or dendrites. This counterintuitive difference compared to anesthesia the reduction of the generalized tonic inhibition of cell bodies (*damping* + *slopeo_damping*) is not altering crucial integration steps (see **Fig. 2c**). Additionally, sedation in sleep is indicated by a low pass filtering, but still showing freely behaving lognormal processing (**Fig. 3b**). Localization and dissociation are indicated by a power law (**Fig. 3c**). Interestingly Ribeiro *et al.* (2010; Figure 4a-c)[34] shows the persistence of lognormal avalanche fitting of slow waves and waking state, as well as the transform to power law tail mediated by anaesthetics.

**Regarding overtones**, Laxpati *et al.* (2014)[17] show a detailed analysis of overtones production in rodent hippocampus by applying short pulsed inputs of 7, 17 and 35 Hz. We repeated this experimental procedure in the non-local model for 7 Hz (**Fig. 3d**) using short pulsed input (1 ms). It was shown that the non-local model also produces overtones similar to that of Laxpati *et al.* (2014)[17]. On top of that, we could not only reproduce the overtones, but also show that peak input only generates overtones, whereas sinusoid input decreases overtones, or totally prohibits overtones (**Fig S2**, compare to Laxpati *et al.* (2014)[17] Figure 7).

**In terms of cortical wave speed**, there is a linear dependence of the speed at which the waves travel and the maximum bandwidth that can be coded in the model (**Figure 3e**). This linear relation collapses close to the maximum speed of 0.5 m/s and the maximum frequency that can be generated by the sampling rate of the model. Close to 500 Hz and an NI of >2.6 the speed of the waves converges to 0.5 m/s. A wave speed of cortical waves of 80-107 mm/s at a frequency of ~7 Hz is determined by Lubenov and Siapas 2009 (Figure 4)[45] in the hippocampus. This wave speed goes along well with the simulated wavespeed. Furthermore, a linear relationship of wave speed and frequency is described by Zhang and Jacobs 2015 (Figure 4)[46].

**The performance** of the model is dependent on the length of the signal, the operating bandwidth, the precision of peaks, the sampling rate, information content of a peak in Fourier space and many additional factors. Further, the performance for the non-local model can be estimated theoretically, however, it can simply be estimated empirically. **Figure 4a** shows that 29 peak frequencies from 5 to 250 Hz can be clearly decoded within 3 s. Furthermore, 48 overtones of the input signal are also well decoded. The resolution of the raw signal could be enhanced in frequency space by dividing point-wise through the corresponding moving average (window-size: 30). This example shows that >50 bit of stimuli can be easily coded within 3 s. Furthermore, the half-width of peaks using sine input centers around 0.5 Hz within 3 sec signal length (**Fig. S6**). This suggests that within 3 sec and a bandwidth of 7-500 Hz about 330 bit can be coded. The bit coding can be increased by utilizing a higher sampling rate, bursts and phase coding.

**In non-local processing the size of the model** should not influence the processing power, as each unit is capable of processing the full bandwidth (**Fig. 4b**). In comparison, a hologram is also capable of storing full information in a spot of sufficient size, but quality of reconstructed image increases when the information is distributed over to entire storage medium by a diffusor[47]. **Figure 4b demonstrates that the size of the model** has no influence on the decoding of the input frequencies and there is also no change in artificial frequencies. A 20x20 matrix, as well as a150x150 matrix is equally resolving 12 stimuli without generating artificial frequencies. Thus, it appears as if the overtones are increasing with size, potentially due to decreasing chaotic processing, or the underrepresentation of overtones in the signal. This might indicate increasing precision in processing.

**Distinct importance is given by the baseline activity** that is defined as the activity of cortical areas that organizes after several seconds to minutes without external stimuli[48,49]. The typical baseline activity is around ~8 Hz[50-52]. To date, the origin of baseline activity has not been totally resolved. However, some characteristics go along with it, such as it is not dominant in presence of external stimuli[53,54], as well as during NREM sleep[1,51,55] and it may be related to spontaneous brain activity[50]. In **Figure 4c** we can show that baseline activity appears self-organized in the presence of spontaneous activity and only when no external stimuli are applied. Furthermore, when the energy transmission is decreased to NI 0.1, transferring the model into slow waves state, the alpha activity is decreased to theta wave activity (**Fig. S3**).

**Coherence measures of the cortex** deliver insights into the spatial behaviour of cortical processing. Spontaneous activity alone shows the **coherence pattern** is continuously decreasing. As described by Maier *et al.* 2014 (see Figure 3)[1,56] and Srinath and Ray 2014 (see Figure 1)[31] that coherence declines rapidly with distance after 0-30 Hz (**Fig. 4d**). The model, as well as Srinath and Ray 2014 (see Figure 1)[31] show that the coherence converges at a level of ~0.1 (**Fig. 4d**). It is hypothesized that decoherence is associated with phase coding. However, harmonic input raises coherence at the input frequency. In **Figure S7** we show that a single burst of 100 Hz results in a peak like increase of the coherence at 100 Hz, similar to the stimulus based coherence increase described in Srinath and Ray 2014 (see Figure 1)[31].

**The non-local model** is built on simple rules and complex interference pattern originating from an increasing number of processing units following the simple rules. This concept is called **emergence**. The properties of the system are more than the sum of the properties of each unit. Emergence is also considered as a possible explanation for consciousness[57,58]. Still, emergence is not easy to conceive, determine or measure.

In physics, as well as in biology there are many examples of emergent systems that form complex structures just from interacting modular building blocks. Emergent systems show **critical distributions**, or near critical distribution as shown by the Ising model[33] and within the non-local network (**Fig. 4e**): dynamic structure formation shows a power law distribution maximizing information content still preserving order just below turbulence and chaos[59,60].

**In Figure 4e** we show that the **avalanche distribution of an EEG signal** is not only producing critical distributions, a hallmark of **emergence** and self-organization[12,23] but also shows an extended shift to higher activity by increased number of connectivity, similar to brain observations[33] (**Fig. 4e**). By this, the non-local model reproduces two phenomena associated with observations: brain activity and criticality for best capabilities in information processing.

**Alzheimer's disease** is characterized by a loss of entire processing units[61-63]. The loss of processing units (lesions) is linked to tau and beta-amyloid plaque formation that impair cell function and accelerate cell death[64]. The lesions in the brain appear randomly, but each lesion can grow in size[61-63]. We used this background to build up a very simple A.D. model () that consists of the initialization of a distinct percentage of random lesions that can grow in size. Using this simulation, we investigated a gradual increase of lesions on the coding of the model. At a level of 0.8% of lesions, artificial frequencies remain undetected, overtones reduce from >60 to 32 and matched frequencies decrease from 12 and 6. Only a low level (0.1%) of lesions still permits to resolution of all input signals.

**Schizophrenia** causes correlation problems of signaling between the neurons in synaptic, dendritic, axonal or supporting cell level[65-68]. Correlation problems of signaling between neurons can occur in synaptic, dendritic, axonal or supporting cell levels[65-68] (**Fig. 4g&h**), mirrored and observed in the complex genetics of Schizophrenia[69].

The **benchmarking** approaches of the critical processing indicate that 12 signals are decoded at a **Schizophrenia** level of 0 (so high correlation between groups of neurons). Overtones are 30 and artificial frequencies are 0. **Fig. 4g** demonstrates that with increasing ratio of uncorrelation the number of artificial frequencies increases and the number of resolved frequencies, as well as overtones decreases. At a level of 0.2% of uncorrelation, artificial frequencies increase to 63, whereas overtones and matched frequencies decrease to 42 and 7. Hence, a percentage of 0.02% of uncorrelation still permits to resolve all input signals. Findings regarding schizophrenia evident from the model are that the overtones, as well as the information content that can be decoded by FFT are decreased, as well as the overtones, whereas the artificial signals are increasing.

By analyzing the **beta band dynamics** (**Fig. 4h**) we get insights into the brain functions, such as coding in burst packages, extent of NI and damping, the combination of reference beam and superimposed information, as well as the coding in sine like signals. Those are also parameters that parallel holographic processes and they are captured in the model, including correctly modelling healthy state and beta band decline in Schizophrenia.

**Fast ripples in cortical signals** are associated with stimulation and active information processing[70]. In the simulation a stimulus is composed of a 31 Hz carrier and superimposed information at a bandwidth >400 Hz within a burst period of 50ms. Such a stimulus causes ripple like signals as shown in **Fig. S8**. The simultaneous EEG signal appears ERP like[71,72]. Oritz *et al*. describes similar fast ripple signals[70]. A related video of simulated cortical activity is provided online, https://www.biozentrum.uni-wuerzburg.de/bioinfo/computing/neuro.

**Comparative analysis of in silico and in vitro electrode signals** - indicated in **Fig. S9a,b -** resolve a similarity of the in silico avalanche distribution and the in vitro baseline activity of wt mice and VEPOR++ mice[73]. Stimulation with CCH seems to shape the lognormal distribution, shifting it to higher amplitudes. The goodness of the lognormal fit appears to degrade with time after CCH stimulation. The effect might be more extensive in VEPOR++ mice than in wt mice.

We demonstrate **the effect of undersampling** in **Fig. S10** which is related to active information processing as indicated in **Fig. S8**. Actual coding at high frequency bands >400 Hz might escape attention as integrated signal (EEG) analysis in frequency space suggest coding a low frequencies whereas small scale recording (MEA) implies high frequency coding where information is actually encoded. Therefore, we suggest here high frequency coding which is well supported in literature[74]. In **Fig. S6** we also demonstrate similar half-width as in [74]. Half-width estimate correlate with maximum coding frequency.

**In how far can network biology provide a "theory of everything"?**

We close with a short discussion of a biology-motivated "*theory of everything*" looking at the network properties of any molecular or larger biological object and its environmental interactions. This can be compared with "*theories of everything*" in physics where the focus are the four fundamental forces and their associated symmetries[75] but with implications for our world, the universe and even landscape multiverses as well as fundamental terms and encyclopaedic views in philosophy *(*different definitions and views on *"meaning", "fundamental question", "meaning of life", "theory of everything"*).

In some sense network biology provides a "*theory of everything*" (TOE), as all interactions for a whole ecosystem (or even a planet, a world …) can be studied, stressing all self-organising and living parts while all forces from physics are only implicitly modelled (by the dynamic interactions between or inside the different sub-networks). A "*great question*" certainly shows the ability of the observer to ask fundamental questions and living beings can typically deal with their own foundations (e.g. their own replication, but in more complex organisms also their whole view of the environment) without running into decision problems[76] as decision conflicts are stochastically resolved in such a way that they favour optimal adaptation. Furthermore, living beings have to be rather complex (even irreducibly complex[77]): no shorter programme than their own DNA will reliably reproduce stopping probabilities for the system. This implies (e.g. species within an ecosystem) an even more complex environment. Regarding our *universe* compared to alternatives ("multiverse"[78]) we thus may speculate that this implies a high probability for us (as living beings) to be situated in the most complex of all worlds as this most complex world takes most of all possibilities. Hence our particularly fine-tuned world (favourable for life to exist[79]) is no wonder, but has a high probability by its shear complexity whereas more boring worlds take a small space and have nothing as complex as life.[80]
