## Supplementary material for "Quantifying and modelling non-local information processing of associative brain regions": simulation details

### Tutorial: The “non-local brain” simulation

#### Setting things up

The simulation “non-local brain” is executed in NetLogo using the file ***Netlogo_Non_Local_Information_Processing.nlogo.***

The tutorial explains our program written in *NetLogo* language. It shows versatile applications of our software and how we model different information transmission processes in the brain. Moreover, our software can directly visualize emergent phenomena in neuron populations (eg. in the associative cortex). Such populations are generated defining simple and basic rules for each neuron.

#### Use cases and examples demonstrate the simulation further

##### Interface and examples

This allows to easy modify any of the simulations shown in Fig. 2-4. Follow these steps:

Open ***Netlogo_Non_Local_Information_Processing.nlogo.***

This will display an Interface (Fig. T1)

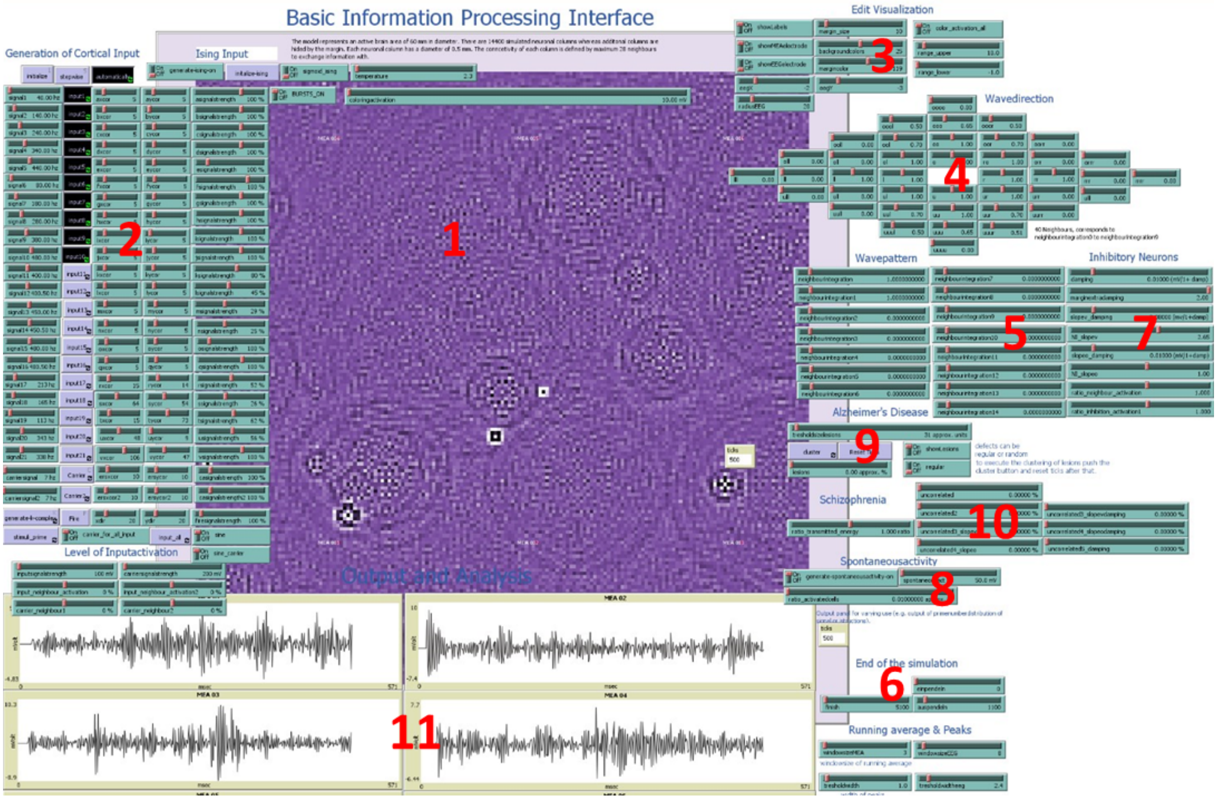


**Figure T1:** Interface of *NetlogoBrainsSlideSimulation*. Elements are: the visualization of activation levels (1), the generation of the rhythmic local field potentials (2), a section to alter the visualization (3), sliders to influence the symmetry of energy transfer (4), a section for adjusting the basic properties of energy transfer, such as damping, number of neighbours, slope (5), parameters to determine the signal length (6), parameters to simulate inhibitory neurons (7), a generator for spontaneous activity (8), simulators for Alzheimer's disease (9) and Schizophrenia (10), as well as the output panel for the depicted signals (11).

The following figures will describe the highlighted feature of the interface seen in Fig. T1 in more detail.

The main screen visualizes the simulation. The top slider (Fig. T1.1) named '*coloringactivation*' literally translates the *activation0 level* in color code. Furthermore, the *tick counter* is displaying the number of *ticks,* or *calculation steps* that symbolize *milliseconds* (red circle).

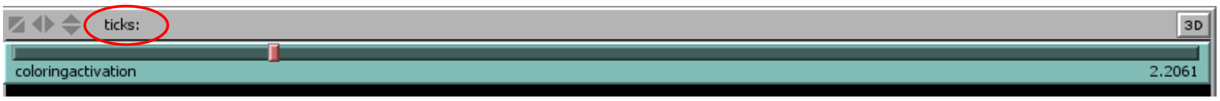


**Figure T1.1:** Slider for coloring activation.

A basic property of the simulation of the cortical area is that energy input can enter the model everywhere, with different energy levels and distinct frequencies. Fig. T1.2 demonstrates the parameters that can be altered to set up and fine tune the input. Aside from this, even more complex signals or chaotic signals can enter the processing layer of neurons. For this purpose an input can manually be set, e.g. to generate phenomena like k-complex.

Fig. T1.2 shows buttons and sliders where you can 'initialize' the simulation and let it run 'stepwise' or 'automatically'. Furthermore, you can define different inputs (*input1 – 12),* with defined frequencies (*signal1 – 12*), signal strengths and set their coordinates (*xcor, ycor*). There is an additional *Carrier signal* and a *Fire* button that allows you to manually set the input. Moreover, *the level of input activation* is changed by sliders that allow you to define the *input activation* of *input1-12, Fire* and the carrier signal.

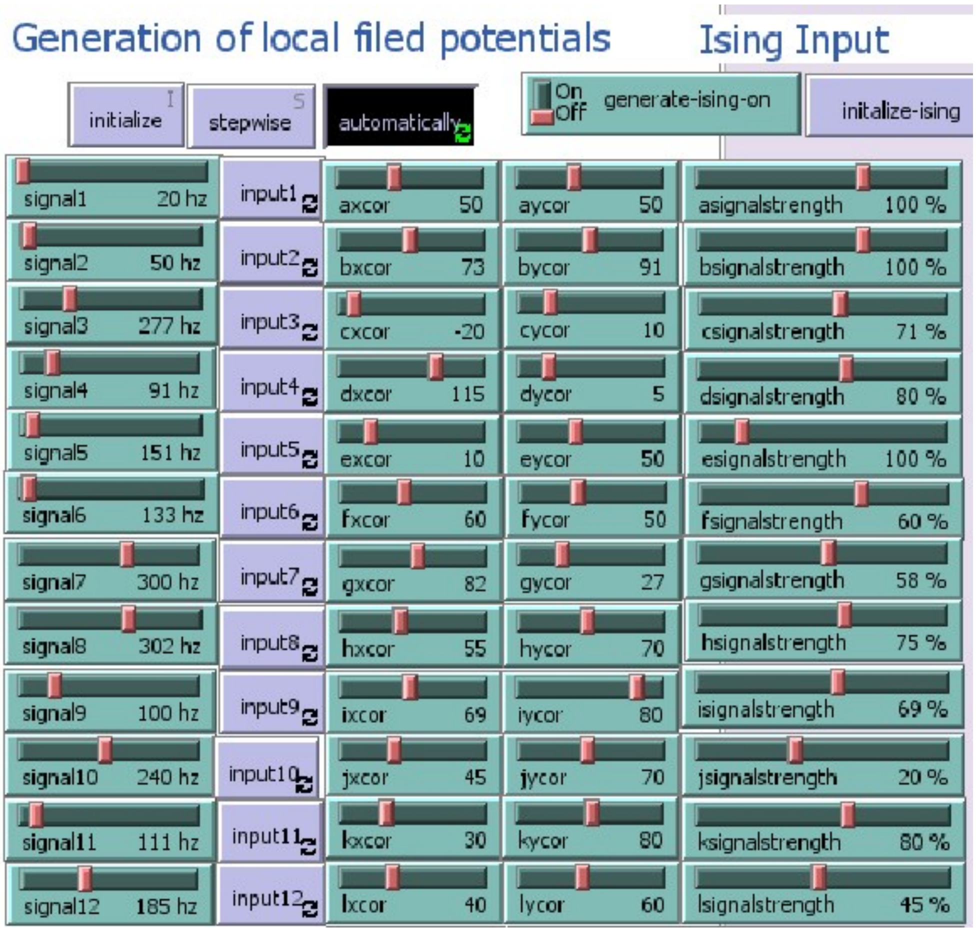


…

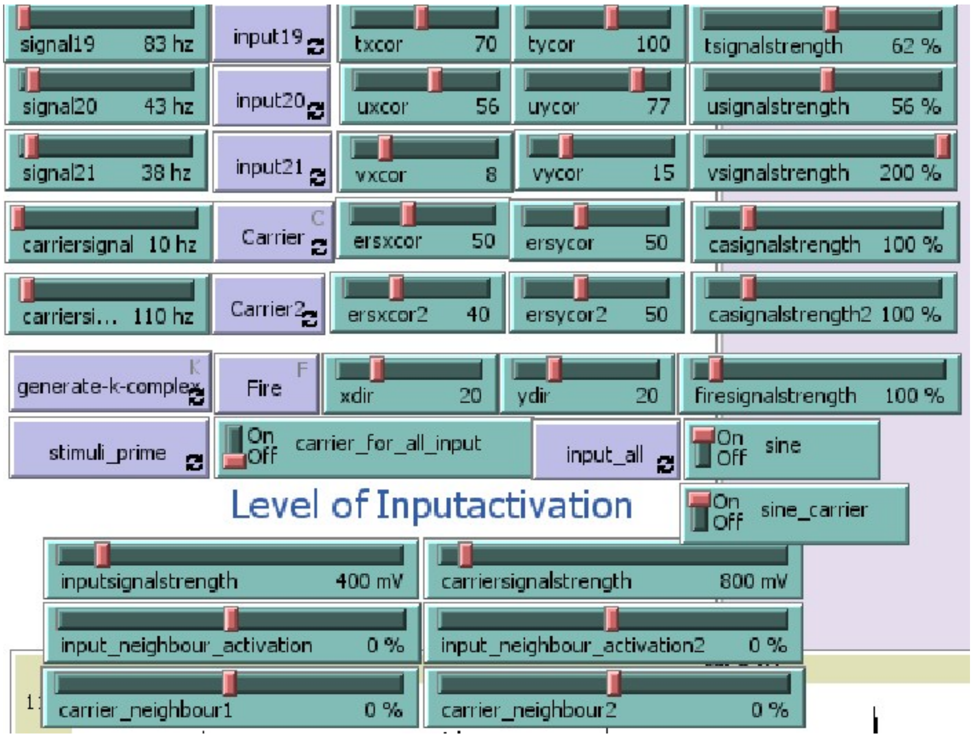


**Figure T1.2:** Demonstration of *sliders* and *switches* for editing the input signals of the simulation. The input signals have a defined strength, a distinct or dynamic localization, are rhythmic or chaotic, can be coded in prime numbers, affect a certain number of neighbours and are generated manually *(Fire)*.

For the purpose of developing a qualitative impression of the interference patterns, the properties of the visualization of the communicating neuronal columns is altered by several 'switches' and 'slider'. In addition, the recording electrodes can be shown and their size and position can be adopted to the object of the investigation.

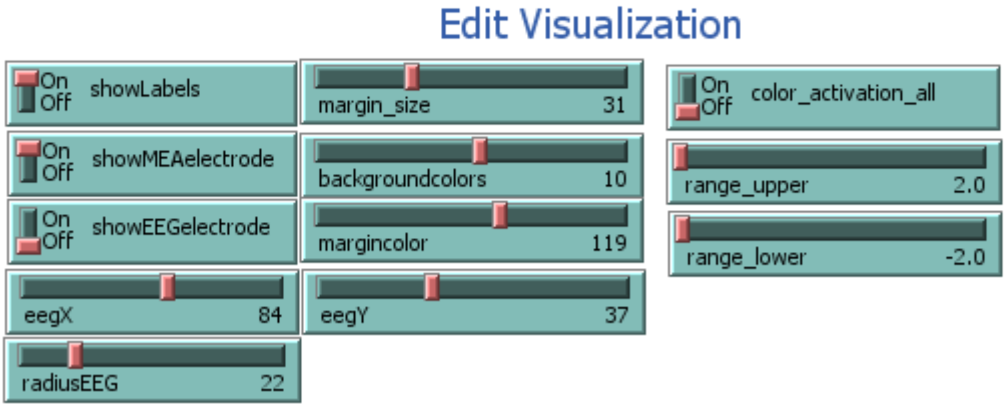


**Figure T1.3:** The graphical appearance is changed with the switches and the sliders, such as *'backgroundcolors'* and '*margincolor*'. An upper and lower range of activation can be selected for focusing distinct energy levels, otherwise the visualization is dynamically adopted to the maxima and minima. In addition, the recording electrodes are visualized on demand and their positions are selected.

Fig. T1.4 shows various sliders that enable the user to alter the *wave direction* in our simulations of neuronal waves. The sliders are ordered according to how the signal is transmitted from a *neuron* or *patch* to its neighbours (e.g. *rr* = 1: information is transmitted to the right-right neighbours multiplied by 1; *or* – upright neighbour; *uul* – lower-lower-left neighbour). By modifying the settings you can verify that the waves are reacting very sensitive to changes. The alteration of the symmetry causes interesting phenomena with complex geometries that not only have regular pattern in space but also in time. Aside from the precise symmetrical information processing gentle symmetry breaking increases drastically the degrees of freedom of the simulation and offers new ways to analyse a systems behaviour facing external world information.

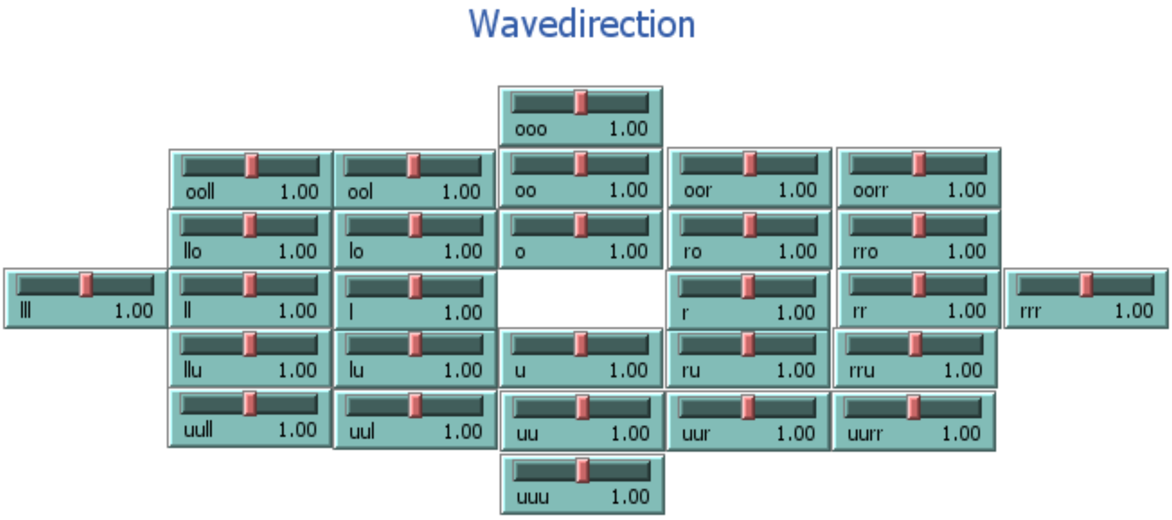


**Figure T1.4:** Alter the symmetry of energy transfer for your purposes. The sliders are in a retinotopic order related to the neighbours of neurons that are exited.

Furthermore, Fig. T1.5 presents *sliders* that offer you options to alter the properties of energy transfer between the patches. In order to simulate transmission of energy between the patches the neighbours are ordered radial symmetric around one patch. *Neighbourintergration* determines the ratio of energy a neighbour contributes with its own activation to the activation of the central patch. *Neighbourintegration* is the ratio of energy transfer of the 4 next neighbours, whereas *neighbourintegration1* is the ratio of the 4 next-next neighbours and *neighboursintegration2* of the 4 next-next-next neighbours and so on (see Fig. 4e). Overall 60 neighbours can be set to contribute to the calculation of the present slope and the corresponding activation level. In this way the activation of the patches is calculated step by step each tick. Similar to an increase of *neighbourintegration* (*NI_slopev*) an increased connectivity can raise the speed of waves (Fig. 3e) and frequency coding potential (Fig. 4a). According to the computational simulation, this will increase the integrational steps to compute and by this running time.

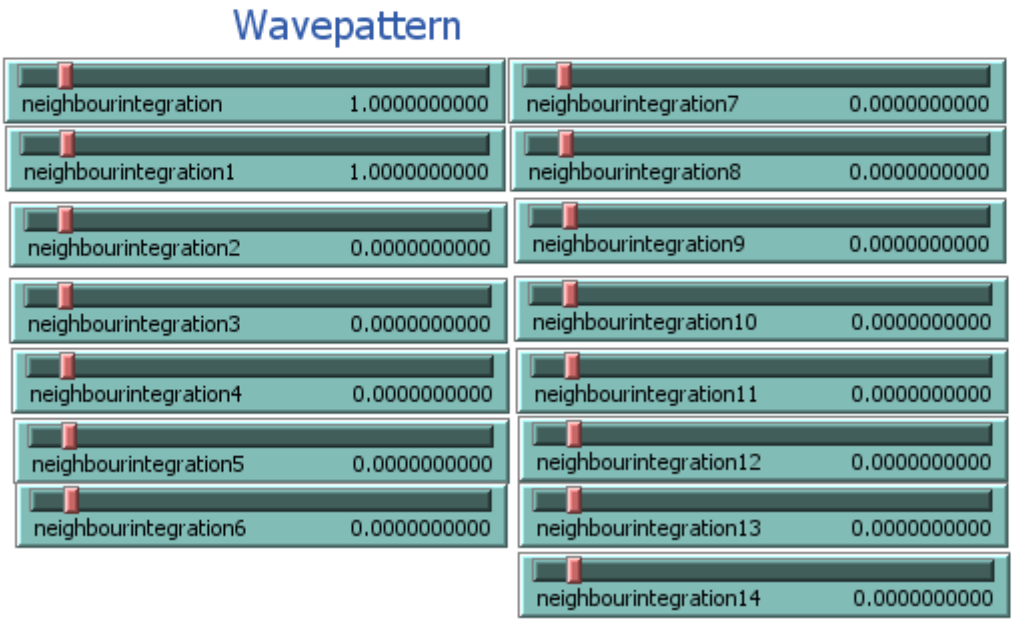


**Figure T1.5:** The properties of the energy transfer can be altered by adjusting the energy transfer between the patches (*neighbourintegration*). An altered energy transfer in time and space will influence the appearance of the interference pattern of the waves the model generates. The number of neighbours contributing to calculation of activation affects the speed of the waves, the wave pattern and the frequency space.

The creation of an interference pattern of different input frequencies in the wave-like brain model urges time. As the model is simulating non-local information processing the supply of each patch with the entire input information urges time that is dependent on the wavespeed and the size of the model. Moreover, the *fft* is performing better with increasing length of signal. Hence, there is a minimum period a model runs to enable investigation of the phenomena of interference. Furthermore, there is a limitation in length of signal due to the processing in short data packages. Smaller data packages with similar bandwidth as longer data packages can process more bits in the same period. Therefore, the regulation of the processing period, as well as the period of processing without input at the beginning and the end of the simulation is crucial (Fig. T1.6).

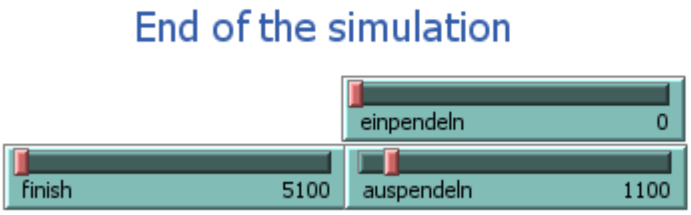


**Figure T1.6:** The signal length, as well as the start of energy input and end of energy input can be determined with the sliders shown.

Fig. T1.7 shows options to implement the balance of excitatory and inhibitory neurons to the model, whereas excitatory integration of neighbour energy is balanced by inhibitory neurons, as otherwise the energy is constantly increasing and the model collapses. There are the sliders *damping (1), marginextradamping (2), slopev_damping (3), NI_slopev (4), slopeo_damping (5), NI_slopeo (6), ratio_neighbour_activation (7), ratio_inhibition_activation1 (8).* According to Fig. 2c and the mathematical description of the model, there are several steps in the main algorithm that can be influenced. Most important is the integration of the neighbouring energy that is set here to 2.65 (*NI_slopev*). A value >=2.7 will generate an exponential increase of energy and frequency and causes the processing to collapse. With increasing number of neighbours, that value has to be adjusted. However, close to the maximum of *NI_slopev* the generation of frequencies close to the Nyquist frequency are permitted. *Slopev_damping* has the reciprocal effect as *NI_slopev*. This is also valid for *NI_slopeo* and *slopo_damping*.

The inhibition of *slopeo* is a damping of integrated *slopev* and damping is the inhibition of integrated *slopeo*, as described in Fig. 2c. The regulation of *slopev* is variable, although a *NI_slopeo* > 1 accompanied by a *slopeo_damping* of 1, or a damping < 0 both result in amplification of energy and system collapse. Those described system collapses that a linked to exceeding distinct borders of ratios of inhibitory and excitatory neurons resemble the signalling pattern of epilepsy (see Fig. S5).

Frequency coding is very stable. Even changing the balance between excitatory and inhibitory neurons can still ensure a stable frequency processing. Most of those modulatory function described in Fig. T1.7 are constantly convoluting the processed function. As long as the ratios of convoluting functions stay constant during a coding period, the frequency coded signals are equally modulated and the relations of energy change stay the same. Yet, a time dependent change of the modulatory function can alter coding packages differently and shift the frequency.

Required are then some further changes, as the decrease of the *NI_slopev* functions acts like a lowpass filter (Fig. 3a). *Decreasing the ratio of ratio_neighbour_activation and ratio_inhibition_activation1*, simulates the increase of inhibitory neuron activity and likewise increases gamma firing (Fig. S4).

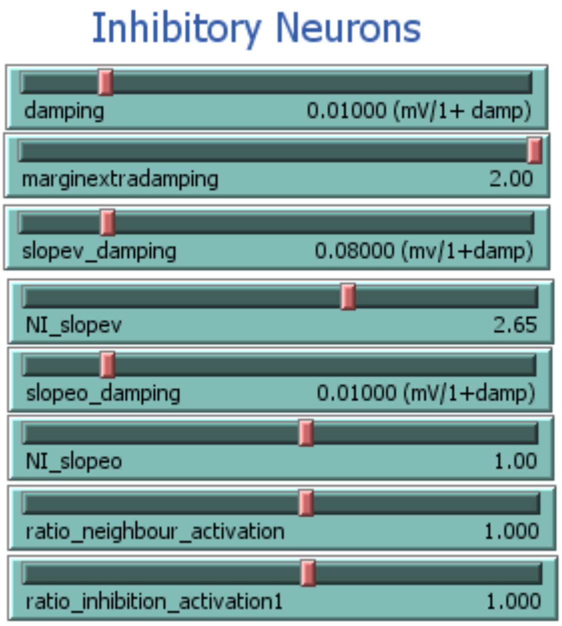


**Figure T1.7:** The modulation of the balance of excitatory and inhibitory neurons is crucial for describing different system states of the brain, like slow wave sleep, waking and active processing. Furthermore, disease like Schizophrenia, Epilepsy Autism may have their origin in intervention of those balances.

Like inhibition, spontaneous activity of neurons is also a complex phenomenon. Spontaneous activity is distributed across the cortex and appears noisy. However, the role of spontaneous activity is not fully understood[1]. In this model you can implement this finding by activating the button *spontaneousactivity* and define the ratio of spontaneous activated cells *(ratio_activatedcells*) and the extent of noise that they are generating (*spontaneousact*). A functional example is shown in Fig. T1.8. Spontaneous activity is applied for analysing the baseline (Fig. 4c & S3).

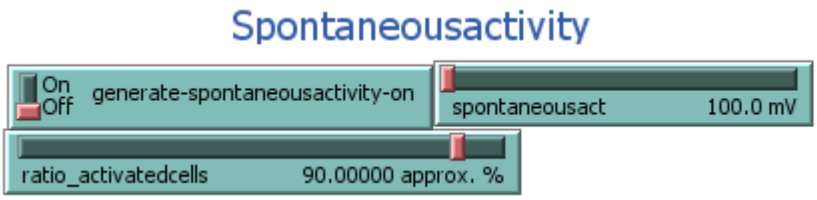


**Figure T1.8**: Generate spontaneous activity of neurons within a neuronal column. Basically, the implementation of spontaneous activity mainly generates different kinds of noise or full randomness, whereas a rhythmic spontaneous activity should not be excluded. Furthermore, spontaneous activity is defined by the ratio of activated cells each processing step and the level of activation.

#### Modelling transmission failure and different diseases

The model allows to simulate different types of transmission failure, mimicking some key features in Alzheimer's Disease and Schizophrenia. The simulator for Alzheimer's Disease (Fig.

T1.9) is based on an algorithm that is setting random neurons to death (inactivating random neurons) – called lesions (*defects* in Fig. T1.9). These lesions can grow by increasing the *threshold2*. The algorithm is built according to description of the development of lesion during the different stadia of Alzheimer's Disease [2,3]. The ratio *of defects* should be set up before initializing the model. After initialization the lesion can be extended by activation the *cluster* button and increase the *treshold2* slider. Before starting the simulation you should push *Reset Ticks.* With the *switches showLesions* and *regular* one can decide to show defects or set them in a regular order. The influence if lesions in coding is shown in Fig. 4f.

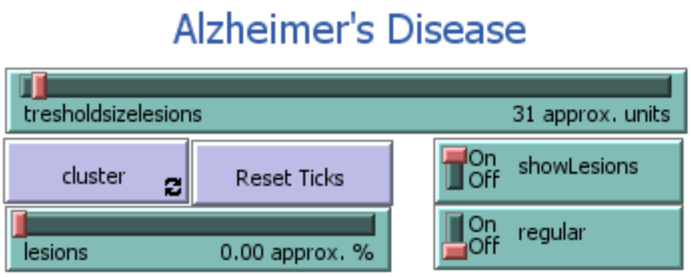


**Figure T1.9:** Simulating stochastic failure of neurons (Alzheimer´s disease).

The Schizophrenia simulation focuses in its basic version on the correlation problems of neurons, that maybe due to a deceleration of dendritic and axonal signalling because of a lack of myelin code, or other cytoskeleton impairments[4-7] . The correlation problems of signalling between the neurons can occur in synapse, dendritic, axonal or supporting cell level[3,8-10] and is related to complex genetics[11]. Nonetheless, in electrode recording, especially EEG recording, we can find an altered complexity of the signalling pattern, indicating that the balance of Glutamate and GABA is shifted, as well as the relation of beta and delta band is declined[3,8,9]. The Schizophrenia simulation is realized by increasing the ratio of *uncorrelated* (Fig. T1.10). The correlation problems randomly affect a distinct percentage of column in the simulation. Similar to the various effects of *neighbourintegration* and its relation to damping, correlation problems can occur on different integration levels of the algorithm. In this way the transmission of the activation1 of the neighbours (*uncorrelated, uncorrelated2*), the *slope_vector* (*uncorrelated3_slopev*), the *slope_old* (*uncorrelated4_slopeo*), or the internal energy duplication (*uncorrelated1*) can be affected. The signals transmission can be totally impaired (*uncorrelated*), or randomly down, as well as upregulated to a distinct range of percentage (*ratio_transmitted_energy*). Besides, the excitatory influence, the inhibition can also be influenced (*uncorrelated3_slopevdamping, uncorrelated4_slopeoddamping,* *uncorrelated5_damping*). This displays a variety of possibilities how a Schizophrenia model can influence the non-local cortex simulation, as displayed in Fig. T1.10. It will be investigated how correlation problems affect the signal processing (Fig. 4g) and which integration level is related to phenomena, such as the beta band decline[12-17](Fig. 4h). Furthermore, uncorrelated signal transmission at various integration levels of the neuronal circuit shown in Fig. 2c, could support the understanding of the complexity of Schizophrenia.

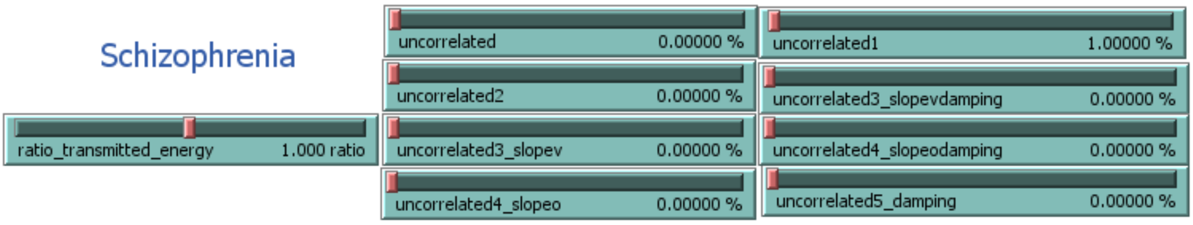


**Figure T1.10:** Low correlation of signal transmission to neighbour neurons: simulating Schizophrenia.

#### Output

Output of the simulation can simulate different detection methods. Thus, the output window (see Fig. T1.11) contains 6 *plots* for displaying *MEA* (multiple electrode array) read-out waves and 6 *plots* for EEG waves (4 not shown). 6 *MEA* electrodes are implemented to compare time, amplitude and phase differences of the measured output. The 6 *EEG* windows allow comparison of EEG recordings from several locations. The slider *finish* determines the number of ticks after which the simulation stops.

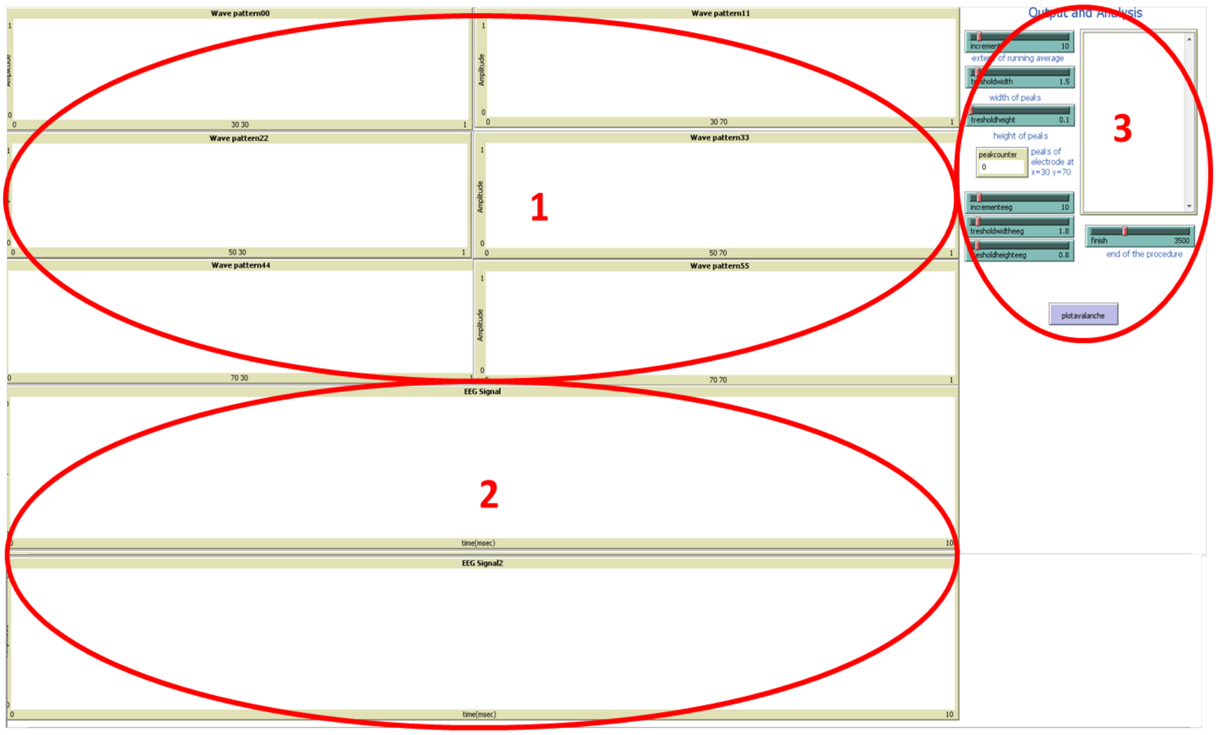

**Figure T1.11:** Output according to MEA recordings (1) and EEG readings (2). Furthermore, it can be adjusted how long the simulation will run (3).

### Analysis Examples

#### Matlab Analysis

After running the simulation for 4100 *ticks* (time steps) the information of the plots - *x-value* and *y-values* - are stored in 12 *csv-files* in the *PackageNetlogo folder*, named: *wavepattern00.csv, wavepattern11.csv, wavepattern22.csv, wavepattern33.csv, wavepattern44.csv, wavepattern55.csv and eegsignbal1-6.csv.* There are example output files in the *PackageNetlogo/Output* folder. With the Matlab files *avalancheanalysis.m*, *fft_peak_analysis.m*, *short-time_fourier-transform.m* and *coherence_analysis.m* several basic analysis used in the paper can be executed.

#### Wave interference

The output window shown in Fig. T1 is giving qualitative evidence of the processing of the model. In the example in Fig. T2 there are 2 external (red stars) and 3 internal stimuli (blue stars) represented and their interference pattern after 20, 50, 100, 150, 200 and 250 ms. Stimuli from the different sensory organs can enter the model at any location of the model. The internal and external stimuli are copied, statistically processed, compared to existing information and integrated. As a consequence, the entered information is distributed over the whole model, which makes it available at any column after a distinct time period. External and internal stimuli generate a complex interference pattern and if not indexed those stimuli are indistinguishable on the level of non-local information processing. It is not clear whether information is lost, modulated or translated in the model, however the bordering conditions and the damping have big influence on maintenance of detecting frequency coded input.

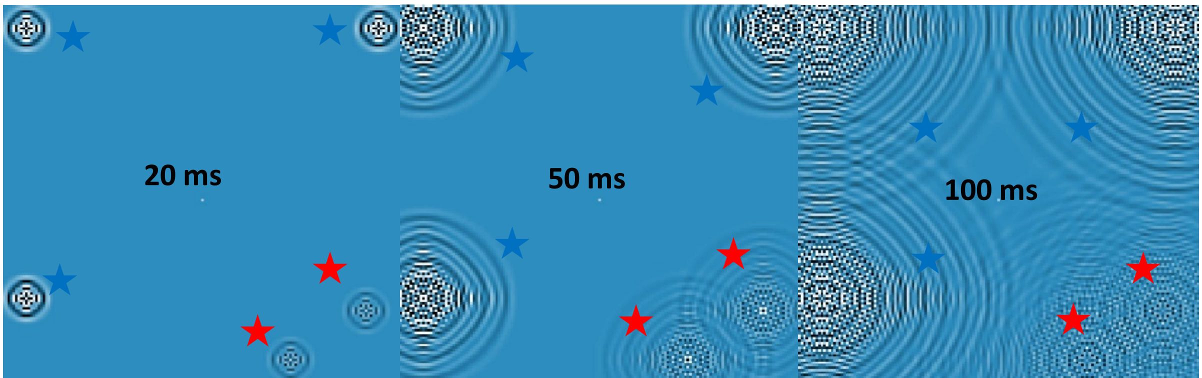

**Figure T2**: Wave pattern is formed by symmetrical interaction of neighbours. Red stars indicate external input, while blue stars indicate internal input. The pictures are made at 20, 50, 100, 150, 200 and 250 ms after stimulus onset. Processing of information is time dependent. Hence, the information spreads over the whole model. Fully integrated information can be depicted from each location. The information can be coded in harmonics and phase by means of the complex interference pattern. With a FFT the input information can be decoded. The frequency coded is dependent on, e.g. the definition of sampling rate, wave speed, wavelength, says space and time.

#### Frequency Analysis

For the functional analysis of data, run Matlab. After that, go to the *PackageNetlogo* folder. Type in the Matlab command line: *fft_peak_analysis_sub.m and show_signal_sub.m.* This will start a Matlab script that runs a Perl parser that compiles the *csv*-files and import them to Matlab-variables for the purpose of plotting MEA-plots and EEG-plots with a running average. On top of this, the data are analysed with a Matlab-implemented fourier transform of each dataset. We show the output of the fft-analysis: The 8 plots generated by the script *fft_peak_analysis_sub.m* are saved in the 8 figures that you can find in the *PackageNetlogo* folder.

Fig. T3 shows an example of 3 plotted original recorded MEA with running average. The MEA consists of 6 electrodes in the model of distinct locations (MEA1: x=30,y=30; MEA2 x=30,y=80; MEA3: x=30, y=90; MEA4: x=80,y=30; MEA5 x=80,y=80; MEA6: x=80, y=90).
The input is peak like, that means that a signal of input 120 mV is applied at distinct time steps. 8 neighbours are contributing to the model with a *NI_slopev* of 1.2. The *marginextradamping* is 0.5 and the input signal strength is 120 mV. The simulation ran 4100 ms, with a period of 3000 ms of constant input. The inputs are 13, 17, 23, 29, 31, 37, 41, 43, 47, 53, 59 and 61 Hz. The input locations are random. The presented signal is 1000 ms in length. Fig. T3 gives a

qualitative overview of the MEA recording and indicates differences that occur despite processing unified input.

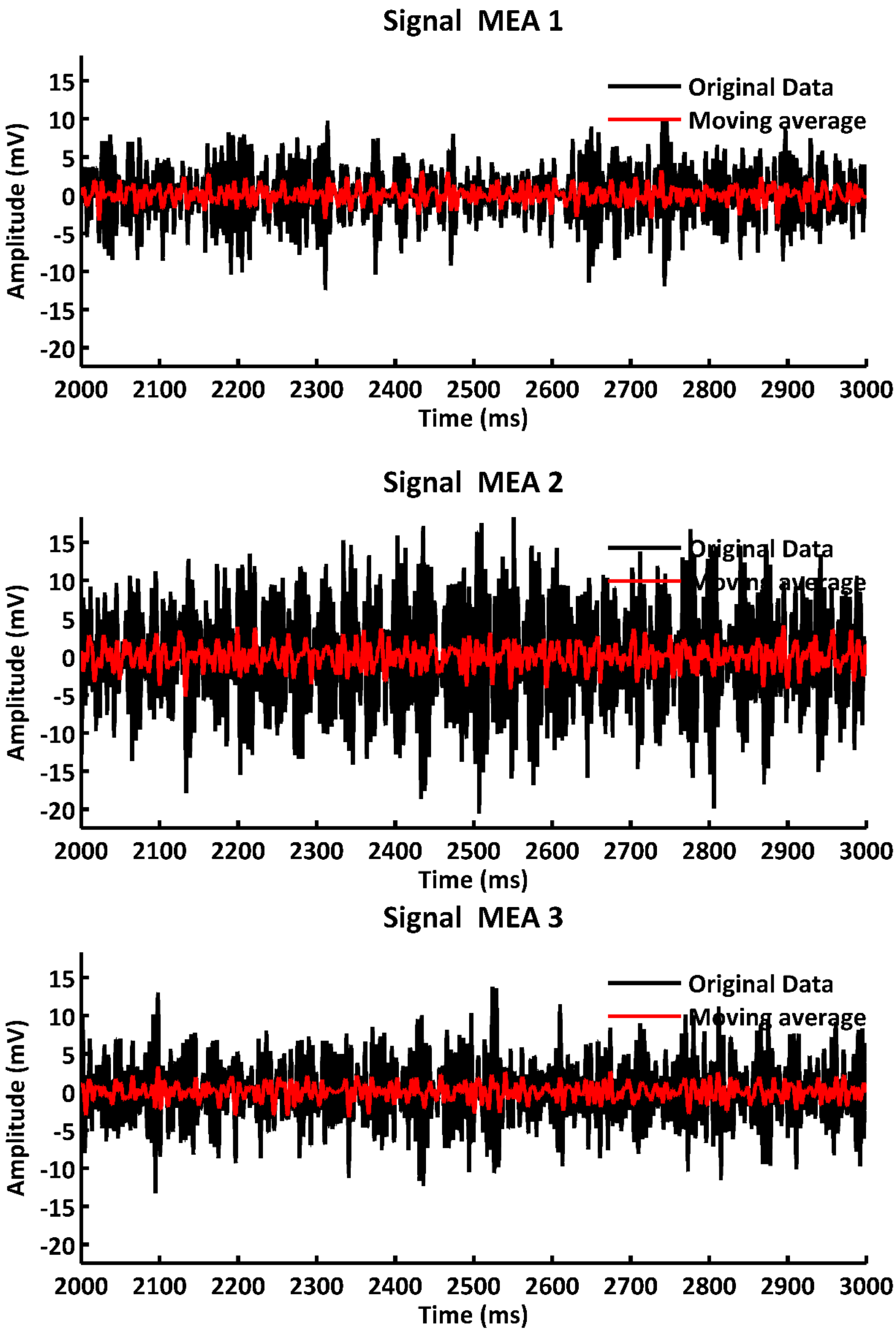


**Figure T3:** 3 MEA-plots of different location of a signal created by the *Netlogo-BrainSlideSimulation* displaying a section of 1000 ms. Each single electrode of the MEA is recording a LFP summarizing input of ~500 µm, corresponding to a neuronal column. The original signal is in black. The running average is marked in red. The wave pattern is created by 12 inputs with peak input of 13, 17, 23, 29, 31, 37, 41, 43, 47, 53, 59, 61 Hz. The location of MEA1 is x = 30 and y = 30, of MEA2 is x = 30 and y = 80, and of MEA3 is x = 30 and y = 130.

Analog to the parameters described above, an EEG pattern is simply generated by integrating all patches in a radius of 10 mm from the EEG location. Central location of the 6 EEG are according to the MEA electrodes (EEG1: x=30,y=30; EEG2 x=30,y=80; EEG3: x=30, y=90; EEG4: x=80,y=30; EEG5 x=80,y=80; EEG6: x=80, y=90). In Fig. T4 examples of the EEG recording are presented and the moving average is added in red.

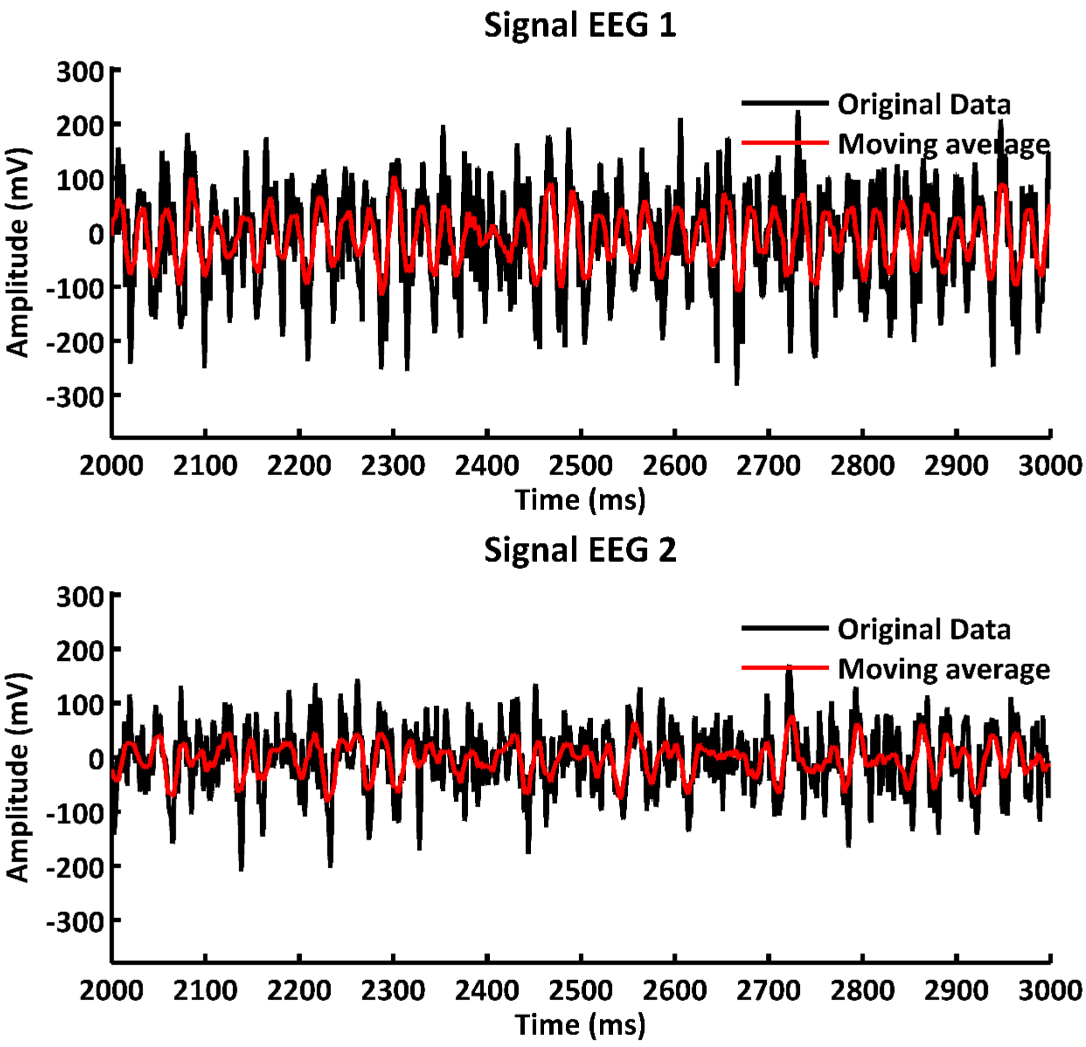


**Figure T4:** 3 EEG-plots of different location of a signal created by the *NetlogoBrainSlideSimulation* displaying a section of 1000 ms. Each single electrode of the EEG is recording a signal integrated over a radius of 20 mm. The original signal is in black. The running average is marked in red. The wave pattern is created by 12 inputs with peak input of 13, 17, 23, 29, 31, 37, 41, 43, 47, 53, 59, 61 Hz. The location of EEG1 is x = 30 and y = 30 and of EEG3 is x = 30 and y = 130.

Fig. T5 and T6 display the fft-analysis of the MEA- and EEG-signal shown in Fig. T3 and T4. The peak frequencies that match to the 6 different *input periods* are highlighted with red circles. The frequencies of 142, 91, 76, 58, 52 and 34.5 Hz. As we suggest frequency coding of the cortex, which well agrees with the topology of the cortex and the non-local simulation, signals are analysed in frequency space (e.g. Fig. 2b, 4a). Interestingly, Fig. T5 shows not only the decoding of the input frequencies (13, 17, 23, 29, 31, 37, 41, 43, 47, 53, 59, 61 Hz), as indicated by blue crosses, but also the presence of overtones of the input signals (green stars). Overtones appear typically when periodic spike input is applied and disappear in presence of rhythmic sinusoid input, as shown in Fig. 3d and Fig. S2). Further parameters for the generation of the input signal are an *NI_slopev* of 1.2, *damping* of 0.03, *mdaming* of 0.5, synchronous processing, an input strength of 120 mV and a signal length of 3000 ms. In Fig. T5 only one electrode is presented of an multi electrode array of 6 electrodes, all measuring activity of a single processing unit.

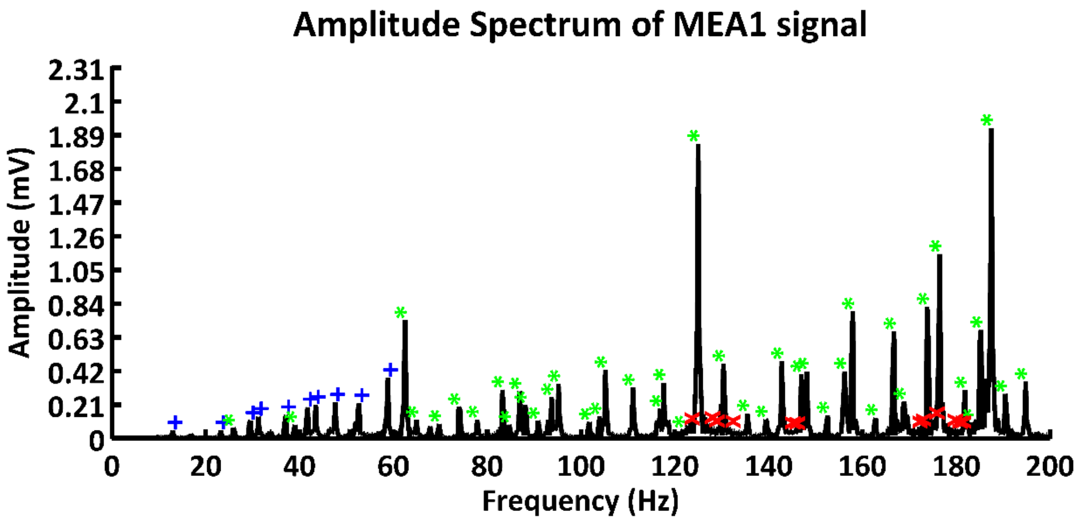


**Figure T5**: The example shows the decoding of an electrode signal of 12 input frequencies (blue) with 17 overtones (green), that appear due to the underlying morphology of the model. Red crosses indicate 4 artificial frequencies, or noise that cannot be differed from the real signal. The parameters are input frequencies of 13, 17, 23, 29, 31, 37, 41, 43, 47, 53, 59 and 61 Hz with an input strength of 120 mV, *NI_slopev* of 1.2, *damping* of 0.03, *mdaming* of 0.5 and a signal length of 3000 ms.

The parameters for the EEG signal are the same for the electrode recordings. The EEG signal is generated by integration of neuronal columns within an area of 2 mm (electrode radius). In Fig. T6 there is also the decoding of input signals, however the amplitudes at low frequencies appear much higher than at the high frequency band in contrast to the frequency space shown in Fig. T5. In both figures there is also the appearance of artificial signals (red crosses). This are either real artefacts caused by input or processing, or artefacts that are based on the precision or sensitivity of the peak detection function. The frequency analysis is executed by the script *ftt_peak_analysis_sub.m*.

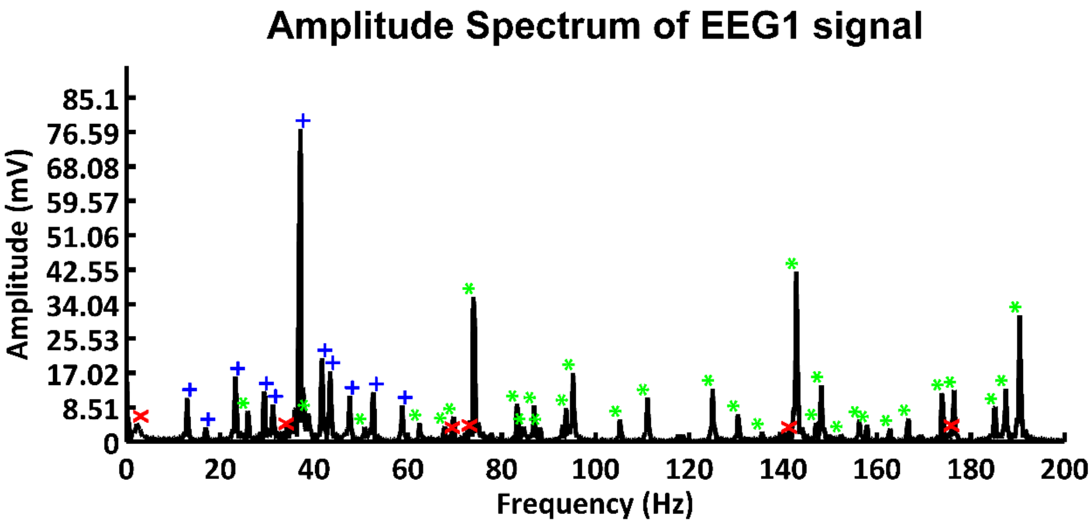


**Figure T6**: The example shows the decoding of an EEG signal of 12 input frequencies (blue) with 17 overtones (green), that appear due to the underlying morphology of the model. Red crosses indicate 4 artificial frequencies, or noise that cannot be differed from the real signal. The parameters are input frequencies of 13, 17, 23, 29, 31, 37, 41, 43, 47, 53, 59 and 61 Hz with an input strength of 120 mV, *NI_slopev* of 1.2, *damping* of 0.03, *mdaming* of 0.5 and a signal length of 3000 ms.

Analog to constant frequency measurements, the short-time Fourier transform (stFT) gives an impression of the dynamic of processing variable stimuli in time. As the cortex is processing stimuli with different length and different phases, or arrival times, the frequency space gets more complex and the resolution in time holds more dynamics and biologically relevant information. In Fig. T7 there is an example of a decoded stationary signal of 12 inputs, transformed in time Fourier space over 4 seconds. It is evident that the constant signal produces a constant frequency behaviour over time. Additionally, the 12 stimuli overtones are again decoded similar shown in Fig. T5 and T6. The stFT is executed by the script *stFT_sub.m*. Related output is also demonstrated in Fig. 4c & h.

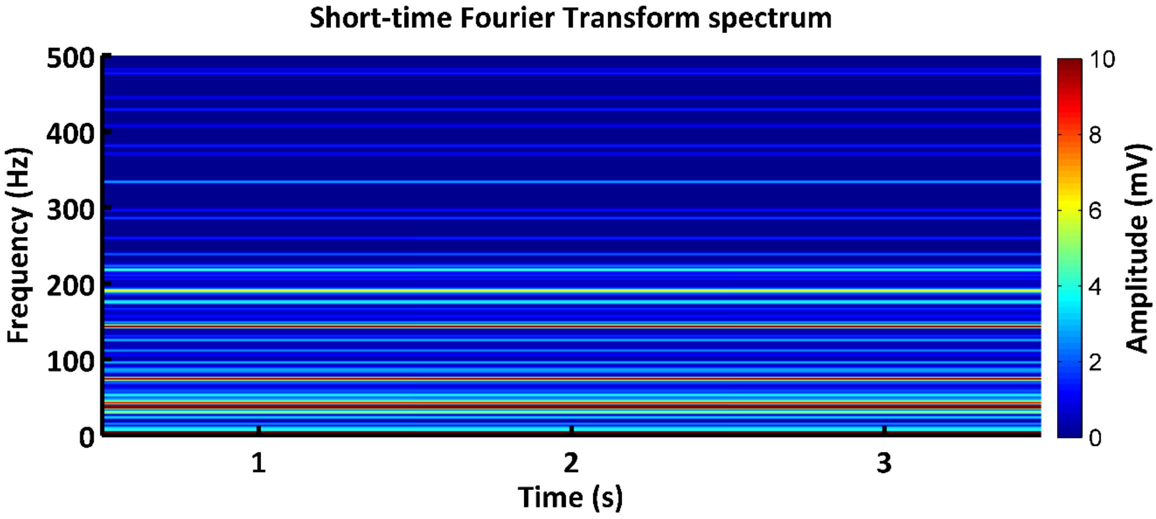


**Figure T7:** Short-time Fourier Transform (stFT) of EEG1 of an integrated signal using the parameter described above. 12 Input frequencies and their overtones can be observed constantly over a period of 3 s. The window size for the stFT is 500 ms and the sampling rate is 1000 Hz. The wave pattern is created by 12 inputs with peak input of 13, 17, 23, 29, 31, 37, 41, 43, 47, 53, 59, 61 Hz. The location of EEG1 is x = 30 and y = 30.

#### Avalanche Analysis

The second anaylsis-script written in Matlab – language is named *avalancheanalysis_sub.m***.** It focuses on the analysis of the peaks of the MEA- and EEG-signal. In order to run the script, open Matlab, switch to the PackageNetlogo/ - folder and type into the *command window: avalancheanalysis_sub.m.* The script will run a peakidentifier and plot the peaks to the signals of the simulation (Fig. T8 & T9). The second part of the script contains a peakcounter and a module to calculate a fit-curve for the plotted avalanche size (peak size) and its related occurrences (P(s)) (Fig. T10 & T11).The avalanche analysis is utilized for a comparison of the *avalanche analysis* of the output of the NetlogoBrainSlideSimulation with a similar analysis of real MEA-data published by Ribeiro[18]. It is to note that the Netlogo model could generate size distributions that are alike size distributions found in freely behaving (FB) animals and anesthesized rats (seeFig. 3b & c). The size distributions help to estimate parameter values such as *NI_slopev*, *slopeo_damping*, *damping*, *ratio of spontaneous activity*, as well as the input type and strength. Hence, the parameters of the model that are adjusted are major driving forces for switching between the brain states.

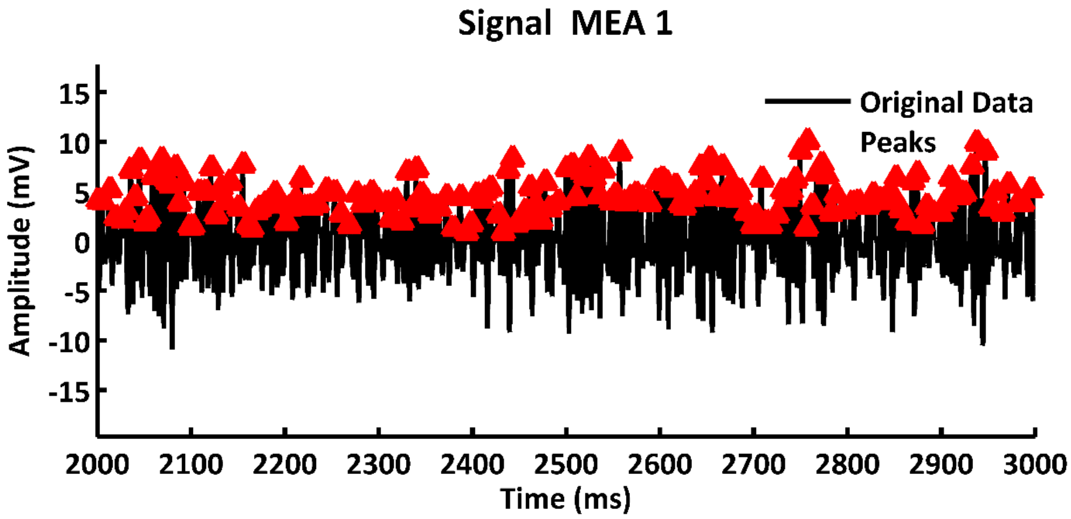


**Figure T8**: MEA-plots with the signal shown in green, running average in red and identified peaks in red triangles. The signal is created by 5 inputs with a frequency of 7, 11, 13, 17, 19 Hz and a carrier signal with a frequency of 29 Hz.

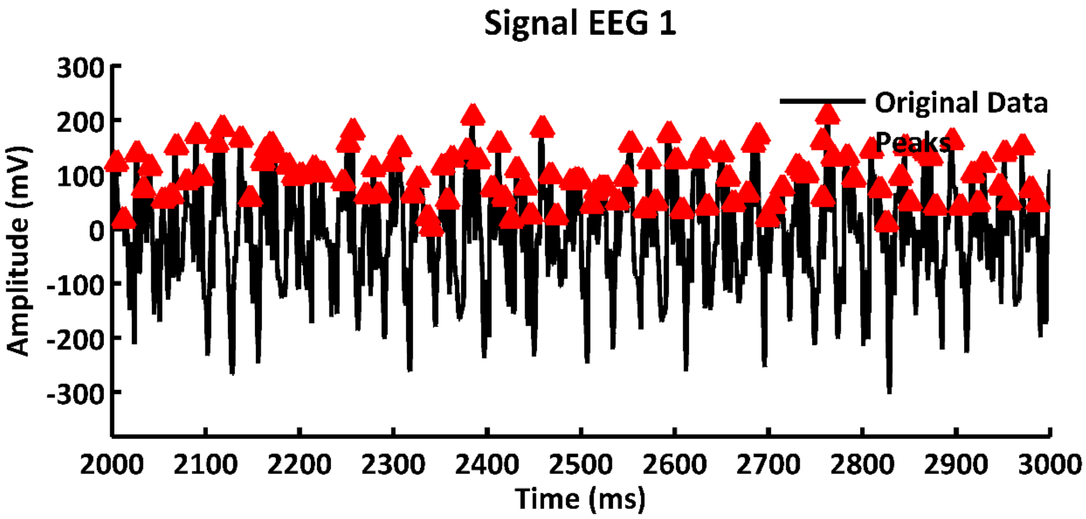


**Figure T9**: EEG-plots with running average in red and peak identifier in red triangles. The signal is created by 5 inputs with frequency of 7, 11, 13, 17, 19 Hz and a carrier signal with a frequency of 29 Hz.

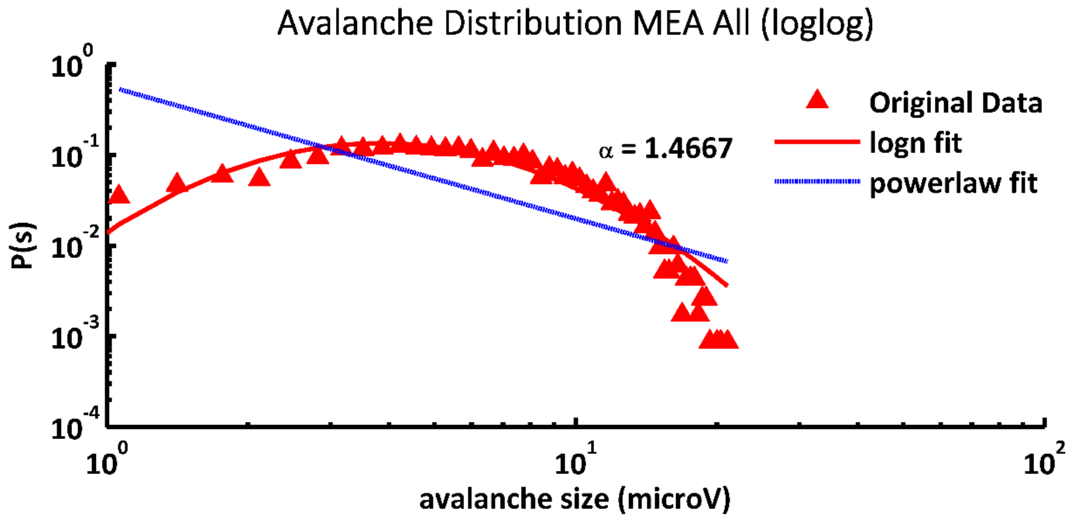


**Figure T10**: The avalanche size distribution of a MEA-signals is indicated in red triangles. The lognormal fit curve is shown as a red line. The power law fit is indicated by a blue line. The signal is created by 5 inputs with frequency of 7, 11, 13, 17, 19 Hz and a carrier signal with a frequency of 29 Hz.

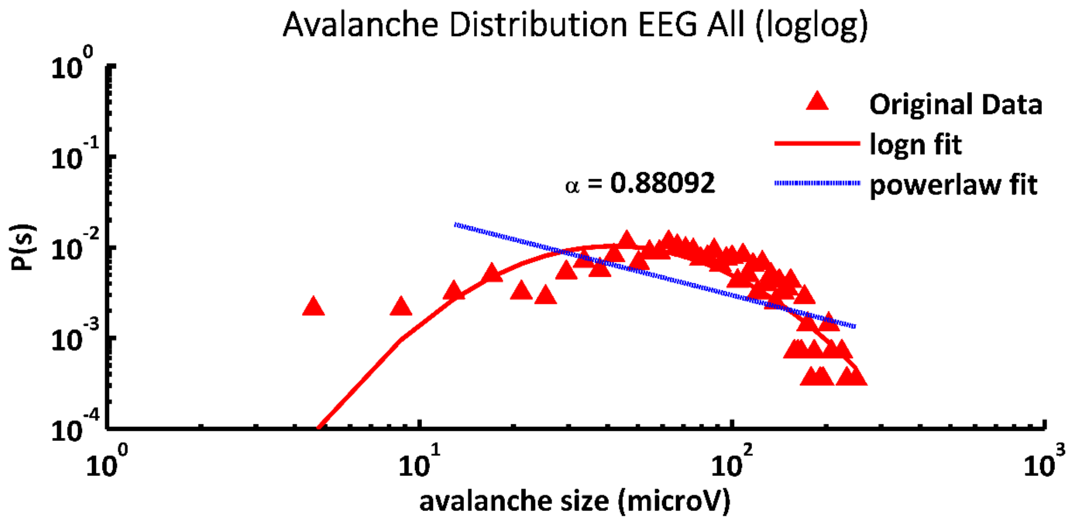

**Figure T11**: The avalanche size distribution of an EEG-signals is indicated in red triangles. The lognormal fit curve is shown as a red line. The power law fit is indicated by a blue line. The signal is created by 5 inputs with frequency of 7, 11, 13, 17, 19 Hz and a carrier signal with a frequency of 29 Hz.

#### Coherence Analysis

The coherence analysis is executed by the script *coherence_sub.m*. The cortex shows distinct coherence profiles when analysed at different frequencies and different tangential, as well as radial distances, as described by Maier et al 2014 (see Fig. 3)[19] and Srinath and Ray 2014 (see Fig. 1)[20]. The coherence is degrading by increasing frequency, as well as with tangential distance (e.g. Fig. 4d). Furthermore, coherence is increasing in the presence of an applied stimuli at similar frequency as the input frequency (e.g. Fig. S7). Tangential distance measurement, as well as frequency measurements are suitable for application in the non-local model. In Fig. T12 we show an example of coherence analysis of a signal created by 1% of spontaneous activated neurons. The input is at random locations each processing step and is 10 mV. The signal is recorded by 6 electrodes at distances of 1 (MEA2), 2 (MEA3), 4 (MEA4), 8 (MEA5) and 16 mm (MEA6), from MEA1. It can be concluded that the coherence is decreasing with distance of electrodes and coherence is declining at high frequencies. However, overall coherence is below 0.6 and is has a maximum at ~50 Hz.

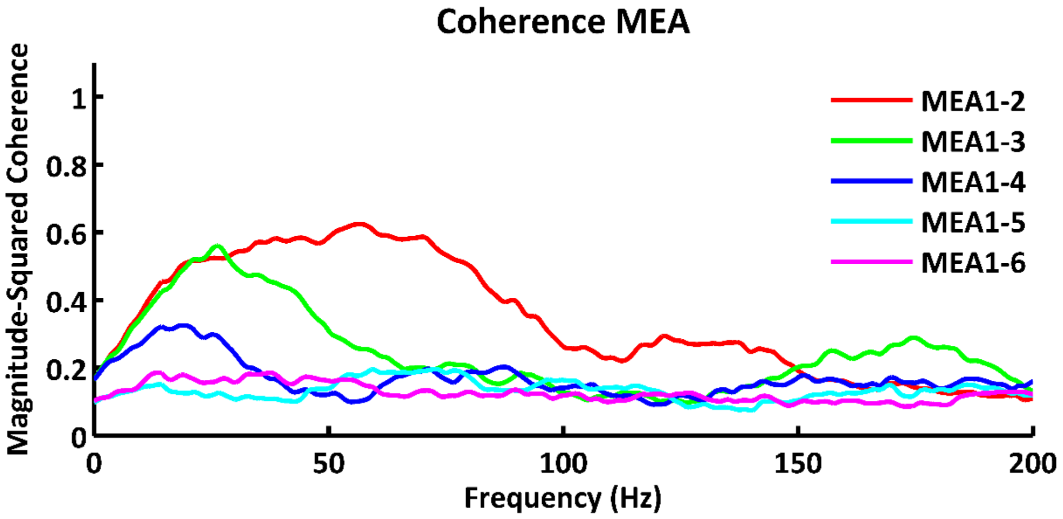


**Figure T12:** Coherence is measured for 6 electrodes. The signals are created by simulating spontaneous activity. Electrodes are placed at distinct distances in linear arrangement. Electrode MEA0 is placed at x = 50 mm and y = 30 mm. The coherence is measured according to the source region MEA0 with distances: 1 mm (MEA1), 2 mm (MEA2), 4 mm (MEA3), 8 mm (MEA4), 16 mm (E5). This figure elucidates the decrease of coherence with rising frequency or distance shown for LFP data by Maier et al 2014 (see Fig. 3)[19], Srinath and Ray 2014 (see Fig. 1)[20]. Parameters: spontaneous activated cells 1% and 10 mV, *NI_slopev* 2.6, *damping* 0.01, *mdamping* 0.5, synchronous, 6000 ms signal length.
